# Computational modelling and near-complete kinetochore tracking reveal how chromosome dynamics during cell division are co-ordinated in space and time

**DOI:** 10.1101/2021.12.16.472953

**Authors:** Jonathan U. Harrison, Onur Sen, Andrew D. McAinsh, Nigel J. Burroughs

## Abstract

Mitotic chromosome segregation is a self-organising process that achieves high fidelity separation of 46 duplicated chromosomes into two daughter cells. Chromosomes must be captured by the microtubule-based spindle, aligned at the spindle equator where they undergo oscillatory motion (metaphase) and then pulled to opposite spindle poles (anaphase). These large and small-scale chromosome movements are driven by kinetochores, multi-protein machines, that link chromosomes to microtubules and generate directional forces. Through automated near-complete tracking of kinetochores at fine spatio-temporal resolution over long timescales, we produce a detailed atlas of kinetochore dynamics throughout metaphase and anaphase in human cells. We develop a hierarchical biophysical model of kinetochore dynamics and fit this model to 4D lattice light sheet experimental data using Bayesian inference. We demonstrate that location in the metaphase plate is the largest factor in the variation in kinetochore dynamics, exceeding the variation between cells, whilst within the spindle there is local spatio-temporal coordination between neighbouring kinetochores of directional switching events, kinetochore-fibre (K-fibre) polymerization/depolymerization state and the segregation of chromosomes. Thus, metaphase oscillations are robust to variation in the mechanical forces throughout the spindle, whilst the spindle environment couples kinetochore dynamics across the plate. Our methods provide a framework for detailed quantification of chromosome dynamics during mitosis in human cells.

## Introduction

The metaphase-anaphase transition is a critical, irreversible step during mitotic cell division, whereby the 46 pairs of duplicated chromosomes (called sister chromatids) are segregated into two daughter cells. During metaphase, chromosomes undergo quasi-periodic saw-toothed oscillations across the spindle equatorial plane (***Skibbens et al., 1993***; ***Wan et al., 2012***; ***Burroughs et al., 2015***). Once the mitotic checkpoint is satisfied, the sister chromatids are pulled towards opposite poles during anaphase (***Musacchio, 2011***). It is crucial that this segregation occurs with high fidelity since errors can cause aneuploidy which is a hallmark of cancer and various developmental disorders (***Gregan et al., 2011***).

The forces necessary for these exquisite chromosome movements are largely driven by interactions between mitotic spindle microtubules and kinetochores multi-protein machines that assemble on each sister chromatid (***Rago and Cheeseman, 2013***). Sister kinetochores are capable of maintaining attachment to both growing and shrinking microtubule bundles (K-fibres) that generate pushing and pulling forces on the chromosomes respectively (***Armond et al., 2015***). Moreover, sister kinetochores are physically connected by centromeric chromatin which behaves as an elastic spring (***Stephens et al., 2013***; ***Harasymiw et al., 2019***) and enables mechanical cues to be transmitted between kinetochores (***Burroughs et al., 2015***; ***Wan et al., 2012***). Such cues control directional switching, giving rise to the metaphase oscillatory dynamic (***Burroughs et al., 2015***). It is this cohesive linkage between sisters that is severed at the onset of anaphase to allow segregation of sisters (***Hauf et al., 2001***), although global phosphorylation states are also important in driving anaphase dynamics (***Su et al., 2016***; ***Vázquez-Novelle et al., 2014***). Sister kinetochores, do not however, operate in isolation in driving chromosome movements; dynamic non-kinetochore spindle microtubules exert forces on chromosome arms and together with arm-tethered molecular motors give rise to a polar ejection force (PEF, (***Ke et al., 2009***)), and microtubule bridging fibers connecting sister K-fibers can slide to push the K-fibre poleward and generate tension between sisters (***Polak et al., 2017***; ***Kajtez et al., 2016***). Within the mitotic spindle, the cross-linked microtubule network produces the highly viscoelastic environment in which chromosomes movements need to be understood (***Shimamoto et al., 2011***).

Chromosomes in human mitotic cells are typically treated as 46 identical objects whilst conclusions are often based on the analysis of (visible/trackable) subsets of sister kinetochores/chromosomes. However, recent reports suggest that there are chromosome specific differences. For instance, chromosomes 1 and 2 have a higher mis-segregation rate than would be expected if all chromosomes behaved identically (***Worrall et al., 2018***), while centromere differences between chromosomes (***Dumont et al., 2020***) and kinetochore size (***Drpic et al., 2018***) have both also been implicated in biasing chromosome segregation errors. A further challenge is that mitotic events occur over multiple time scales, with fast kinetochore-directional switching (on the timescale of seconds) (***Burroughs et al., 2015***), and a slow self organisation dynamic with chromosome congression taking approximately 10-20 mins (***Paul et al., 2009***; ***Auckland et al., 2017***).

Understanding this complex multi-scale mechanical system requires development of quantitative mathematical models that can capture crucial elements of the system’s biophysics and regulatory properties, provide quantitative support for conceptual ideas, and generate testable predictions. Efforts in this direction have been ongoing since the 1980’s with previous work focusing on microscopic models of kinetochore-microtubule attachment (***Hill, 1985***; ***Joglekar and Hunt, 2002***; ***Civelekoglu-Scholey and Cimini, 2014***), on the role of bridging fibres and spindle geometry (***Kajtez et al., 2016***; ***Miles et al., 2021***), and on chromosome congression dynamics to the spindle equator (***Mogilner et al., 2006***; ***Zaytsev and Grishchuk, 2015***; ***Blackwell et al., 2017***). Careful calibration of models to experimental data is crucial to ensure model validity. However, few studies have performed inference of model parameters directly from experimental data. In previous work (***Armond et al., 2015***), we proposed a biophysical model of metaphase oscillations, and fitted the model to 3D kinetochore tracking data from HeLa cells using Bayesian inference, a methodology that propagates uncertainty so parameter confidence can also be determined. The fitted model provided insight into the forces acting on kinetochores across directional switching events, and how sister kinetochores coordinated directional switching (***Armond et al., 2015***; ***Burroughs et al., 2015***).

However, a high degree of heterogeneity in oscillatory dynamics has also been revealed, *i*.*e*. there was substantial variation of the biophysical properties. In particular, the position of the chromosome within the 3D spindle influences mechanical forces with both the polar ejection forces (***Armond et al., 2015***; ***Civelekoglu-Scholey et al., 2013***) and kinetochore swivel (***Smith et al., 2016***) increasing towards the periphery of the metaphase plate. Non-sister kinetochores can also influence each others’ behaviour and exhibit motion correlated to that of their neighbours (***Vladimirou et al., 2013***), hypothesised to be due to cross-linking connections between K-fibres (***Vladimirou et al., 2013***; ***Elting et al., 2017***). To further investigate spatial interactions within the cell and spatiotemporal dynamics through to anaphase, a biophysical model is required that includes the transition from metaphase to anaphase.

In this work we present a metaphase-anaphase model of kinetochore dynamics for human retinal pigment epithelial cells (RPE1); a karyotypically stable, non-transformed cell line. The model incorporates metaphase oscillations, captures the transition to anaphase and the segregation of chromosomes to respective poles. Chromosome movements are driven through force balance between the 4 primary forces acting on a chromosome in mitosis: the K-fibre forces, the PEF, the centromeric spring connecting sisters and drag. We use a stochastic differential equation formulation for the mechanics and a discrete hidden Markov model to model the K-fibre dynamics of the chromatid pair and the transition into anaphase. Using Bayesian inference, specifically a Markov chain Monte Carlo (MCMC) algorithm, we parametrise our biophysical model, providing a powerful tool to interpret/annotate and analyse experimental trajectory data. By combining lattice light sheet microscopy (LLSM (***Chen et al., 2014***)) and endogenous protein labelling, we achieve a high signal-to-noise ratio and temporal resolution whilst ensuring minimal photobleaching and phototoxicity. We demonstrate near-complete 3D tracking of the 46 kinetochore pairs for up to 15 mins. This allows fitting of the metaphase-anaphase model to kinetochore trajectories and characterisation of the biophysical parameters describing chromosome dynamics over the population of chromosomes in human cells.

Our analysis provides a comprehensive atlas of kinetochore dynamics in normal human cells throughout metaphase and anaphase, revealing how spatial positioning defines the mechanical behavior of sister kinetochores such that kinetochore behavior is less variable between cells than previously thought. We show local spatial coordination of directional switches and the timing of anaphase onset. We further demonstrate how dynamics mature as anaphase approaches, caused by a stiffening of the kinetochore-microtubule attachment. Together, these results reveal how interactions between kinetochore pairs can account for the coordination of directional switching events and anaphase onset itself.

## Results

### Near-complete kinetochore tracking through the metaphase-anaphase transition

To obtain insight into chromosome dynamics at the anaphase-metaphase transition, we developed a tracking algorithm that achieves near-complete tracking of all 46 fluorescently labelled kinetochores, using an endogenous label of a kinetochore protein (***Roscioli et al., 2020***). The tracking pipeline is outlined in Supp. Fig. 1 and consists of: deconvolving the 4D movies; detecting candidate spots via an adaptive threshold technique; refining spot locations using a Gaussian mixture model to provide subpixel resolution; fitting a plane to give the metaphase plate as a reference coordinate system; linking detected particles between frames over time to form tracks; and grouping kinetochore sister pairs based on metaphase dynamics. This provides subpixel resolution for the positions of each kinetochore, and allows us to study dynamics of sister kinetochore pairs, rather than simply individual kinetochores.

We performed live-cell imaging of untransformed human RPE1 cells using LLSM (Fig. 1) and generated tracks with our tracking pipeline (details in Methods). Data were collected at a high temporal resolution of 2.05s per *z*-stack over long timescales, typically tens of minutes, from metaphase through to anaphase. A higher time resolution was required than in previous work (***Sen et al., 2021***) (which used 4.7s per *z*-stack) to properly assess fast directional switching dynamics. We obtain partial tracks (in the coordinate system indicated in Fig. 1B) for all 46 sister pairs of kinetochores for the cell shown in Fig. 1A, with 22 of these extending throughout the duration of the movie from metaphase through to anaphase. Across a population of *N* = 58 cells, we obtain an average of 72 [quartiles Q1-Q3:65-77] kinetochores (Fig. 1C) and 29 [24-34] sister kinetochore pairs (Fig. 1D) tracked through at least 50% of the movie.

**Figure 1.**
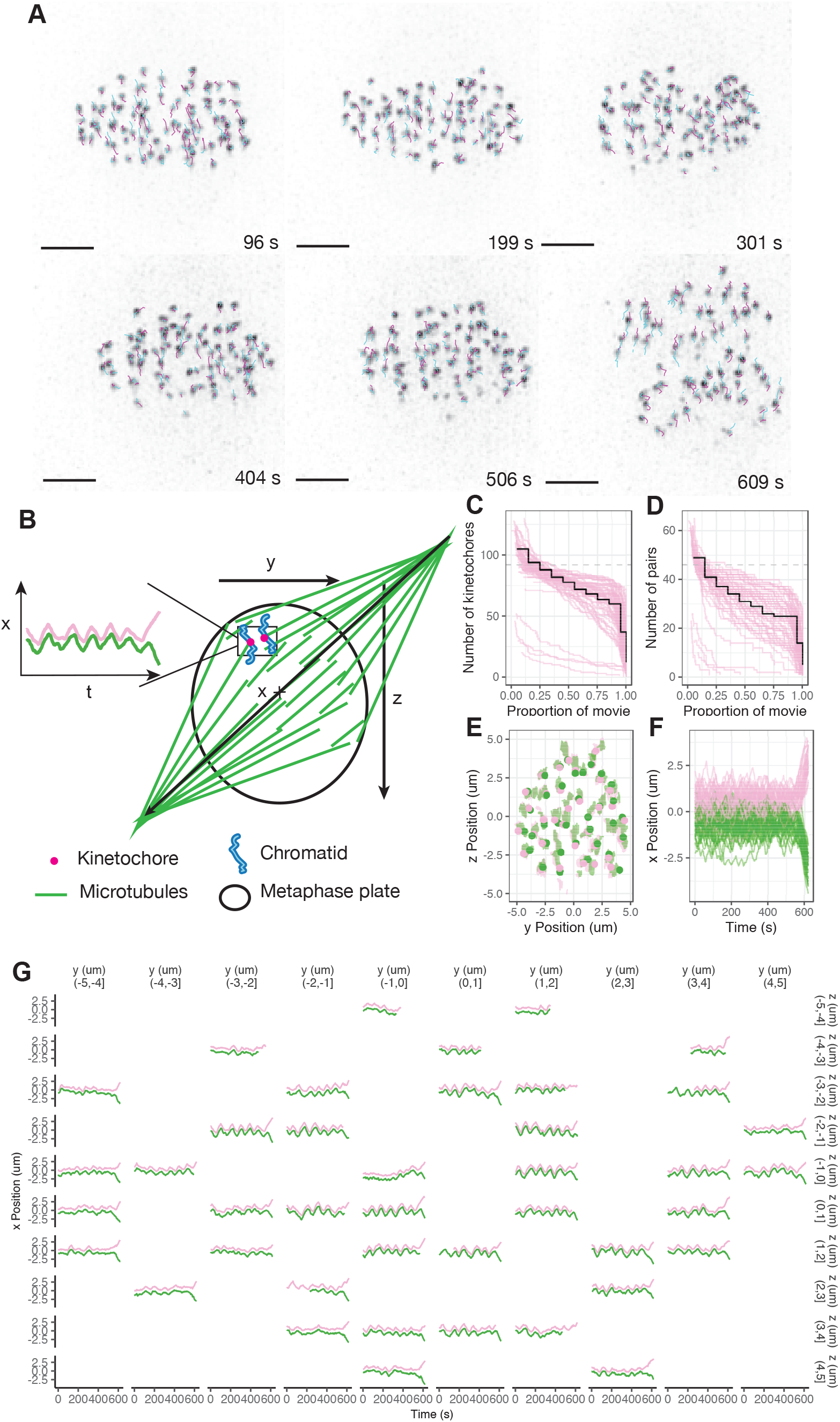
Near-complete tracking of kinetochores through metaphase and anaphase A in human RPE1 cells. (A) Sequence of *z*-projected images showing metaphase in the lead up to anaphase, and the metaphase-anaphase transition in a movie of 619s duration. Dragontails indicate tracked kinetochores in an example cell; cyan (magenta) lines show previous (next) 5 frames of trajectories. Scalebar shows 3um. (B) Schematic showing the coordinate system and metaphase plate. The *x* axes is perpendicular to the metaphase plate, with *y* and *z* axes mutually perpendicular within the metaphase plate. (C) The number of kinetochores tracked through at least a given proportion of the duration of a movie. Magenta lines show individual cells and the black line shows the median over the population of *N* = 58 cells. Dashed grey line indicates 92 kinetochores. (D) The number of kinetochore pairs such that both kinetochores in a pair are tracked for at least a given proportion of the movie. Magenta lines show individual cells and the black line shows the median over the population of *N* = 58 cells. Dashed grey line indicates 46 kinetochore pairs. (E) Positions of kinetochores for the cell in (A) are shown within the metaphase plate during metaphase, with sister kinetochores coloured green and magenta and circles indicating the position at 492 s. Kinetochores have a set position within the metaphase plate with limited diffusion in the plate over time. (F) Tracks for all kinetochore pairs in the cell from (A) showing position relative to the metaphase plate over time perpendicular to the plate. (G) Trajectories of kinetochore pairs over time for the cell from (A) shown with subplots positioned corresponding to their average position within the metaphase plate.

As expected, kinetochores form a metaphase plate (Fig. 1E) and undergo saw-toothed oscillations perpendicular to the metaphase plate (Fig. 1F) before separating in anaphase when kinetochores segregate towards their respective spindle poles (Fig. 1F). Visualising trajectories over time highlights spatial differences between the kinetochore pairs distributed throughout the metaphase plate (Fig. 1G). In particular, kinetochores at the centre of the metaphase plate have greater amplitude of oscillation compared to the edge of the plate (Fig 1G), as expected from previous work in HeLa cells (***Armond et al., 2015***) and Ptk1 cells (***Civelekoglu-Scholey et al., 2013***). Quantitative analysis of variation of the dynamics across the metaphase plate requires a computational model to annotate the trajectories and estimate parameters.

### Metaphase-anaphase model of kinetochore dynamics in human cells

In order to address hypotheses about how chromosome segregation is co-ordinated in space and time, we extended the metaphase dynamics model of ***Armond et al. (2015***) to incorporate the metaphase-anaphase transition. Let 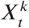 denote the (1D) position of kinetochore sister *k* at time *t* in the *x* direction perpendicular to the metaphase plate. Metaphase dynamics are described by the stochastic differential equations

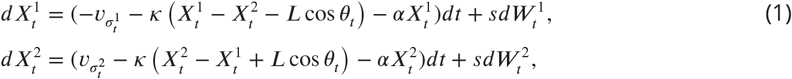

where *θ*_*t*_ is the angle between the normal to the metaphase plate and the vector connecting a sister pair at time *t* (thereby projecting the spring force perpendicular to the metaphase plate), *α* is the polar ejection force parameter, *κ* and *L* are the spring constant and natural length of the centromeric chromatin spring connecting the kinetochore sisters, 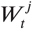 are Weiner processes for sister *j, s* the magnitude of the diffusive noise, *υ*_+_,*υ*_−_ the velocities associated with polymerising (+) and depolymerising (−) microtubule states respectively, and the microtubule states 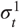 and 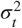 are hidden states taking values in the set

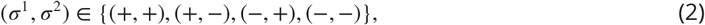

with +/− states denoting polymerising and depolymerising K-fibers respectively. The term 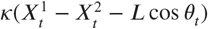 is a Hookean spring force term due to the centromeric chromatin spring connecting a kinetochore pair. The 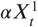 term corresponds to the polar ejection force (PEF) pushing the chromosomes towards the equator. We assume that the PEF is linear in the spring extension near the metaphase plate (***Ke et al., 2009***). The 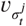 term (taking values *υ*_+_ or *υ*_−_) represents the polymerisation/depolymerisation force from the K-fiber attached to sister *j*, (dependent on the current microtubule state 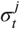). The schematic in Fig. 2A illustrates the forces acting on a kinetochore sister pair, and how these forces depend on the configuration of hidden states (Fig. 2B) at any given time. The stochastic differential equations (1) follow from force balance, dividing through by the unknown drag coefficient as in ***Armond et al. (2015***) (see Appendix 2 for further details); the effect of this is that all terms in eq. (1) have dimensions of speed, and units of the force parameters *α* and *κ* are [*s*^−^1]. We integrate over a time frame, Δ*t*, approximating the system of equations in (1) as a discretised system of equations as follows:

**Figure 2.**
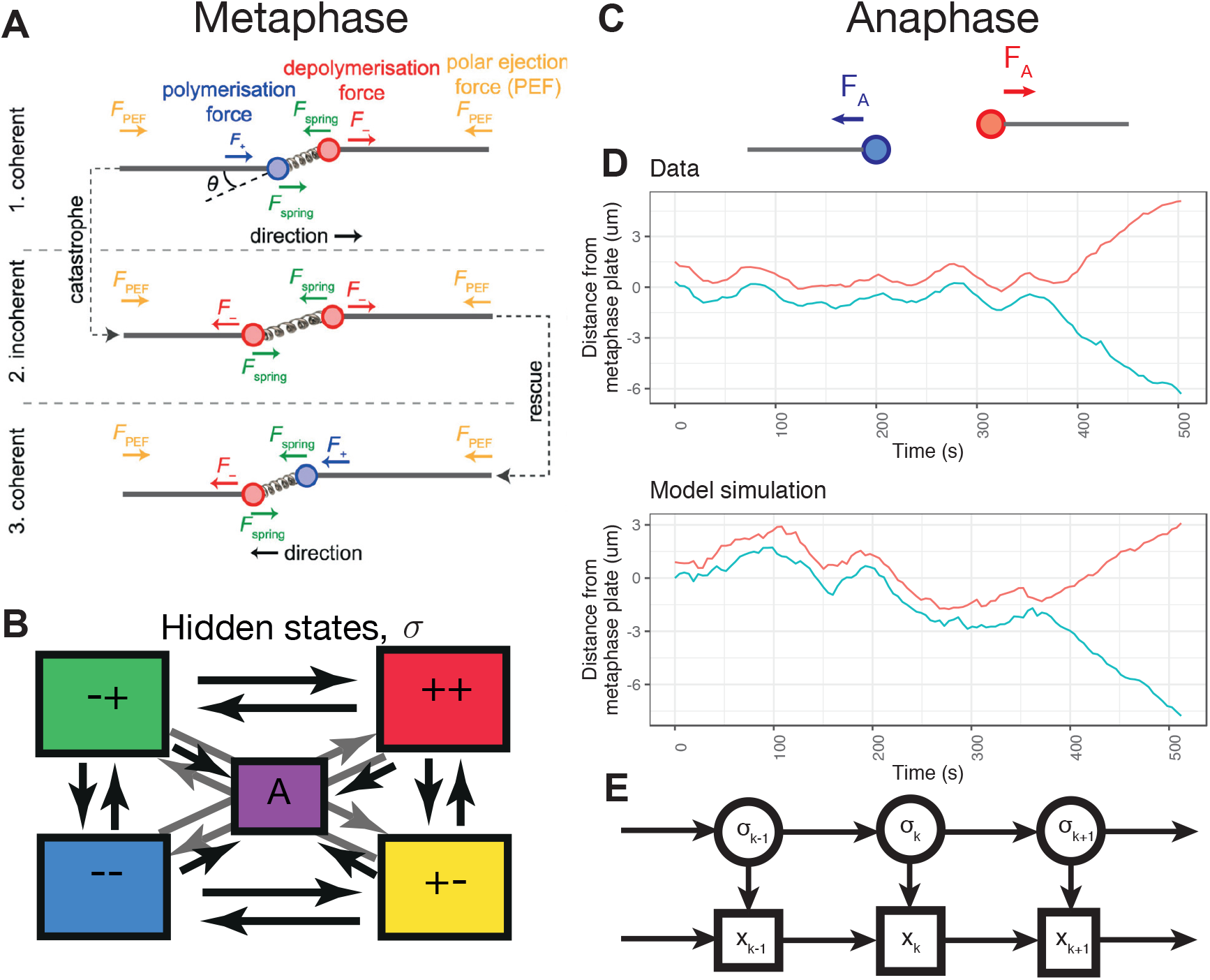
Biophysical model for metaphase-anaphase transition in mitosis in human cells. (A) Schematic of forces on a kinetochore pair in metaphase (inset comparison with zoomed in pair); adapted from ***Armond et al. (2015***). (B) Schematic of hidden states and transitions between hidden states. (c) Schematic of forces on a kinetochore pair in anaphase (inset comparison with zoomed in pair after anaphase onset). (D) Comparison of forward simulation from the model with experimental data for a single pair. (E) Graphical model structure as a state space model, similar to a Hidden Markov model. Here the K-fibre polymerisation states 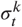 are unobserved hidden variables.

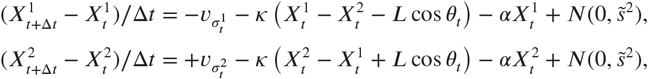

where 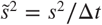.

To describe anaphase, we introduce an additional hidden state, *A*, with dynamics (Fig. 2C) in this state given by

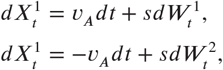

where *υ*_A_ is the velocity in anaphase. We have removed the terms for the spring forces and the polar ejection force (as these are lost at anaphase onset), and allowed the velocity in anaphase to be different to the velocities associated with polymerising or depolymerising microtubule states. The transition to anaphase is controlled by a smooth switch whereby there is a transition probability at time *t* given by,

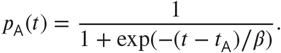

Thus, the transition to anaphase occurs around time *t*_A_ with *β* = Δ*t*/2 determining the range over which switching can occur. We assume the anaphase state *A* is accessible from each of the other states, but transitions back from anaphase to metaphase are not possible, state *A* is absorbing. Thus, once anaphase onset occurs the chromosomes segregate.

Switches between hidden states (see Fig. 2B) occur at each time step according to the time dependent transition matrix (state order as (2), with anaphase state *A* at the end),

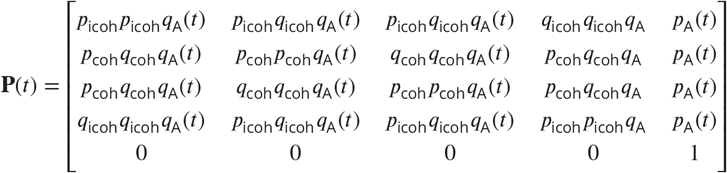

with *q*_coh_ = 1 − *p*_coh_ and *q*_icoh_ = 1 − *p*_icoh_ and *q*_A_(*t*) = 1 − *p*_A_(*t*), where *p*_*coh*_ and *p*_*icoh*_ are the probabilities of a kinetochore remaining in the coherent (sisters move in the same direction), respectively incoherent (sisters move in opposite direction) state over a time interval Δ*t*, and *p*_*A*_ is the probability of transition to the anaphase state, *A*. Simulating from this biophysical model produces trajectories with quasi-periodic oscillations qualitatively similar to observed data (Fig. 2 D). Saw-tooth like oscillations occur when the coherent mean lifetime is larger than the incoherent lifetime.

### Bayesian inference enables automated annotation of microtubule attachment hidden states and biophysical parameter estimation

Automated annotation of kinetochore trajectories is a key tool in the analysis of metaphase dynamics, the transition to anaphase, and inferring potential mechanisms. We take a Bayesian approach to fit the model described in the previous section directly to experimental data. We use MCMC techniques (***Gelman et al., 2014***) to obtain samples from the posterior distribution *P* (*θ* |*x*_1*:T*_) of the model parameters *θ* = (*τ, α, κ, υ*_−_, *υ*_+_, *L, p*_coh_, *p*_icoh_, *υ*_*A*_, *t*_*A*_) given the observed time series data where 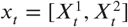 is the observed position of the sisters at time *t*. Here 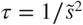 is the precision. Furthermore, we can sample from the distribution of hidden microtubule states, *P* (*σ*_*t*_| *x*_1*:T*_) allowing us to annotate trajectories.

The structure of the model considered here is that of a state space model, very similar to the standard hidden Markov model (HMM) set up, except that we have additional dependencies on the previous observation term, as shown in Fig. 2E. We derive a version of the forward-backward algorithm (***Rabiner, 1989***) that accounts for these additional dependencies, thereby allowing sampling from the distribution of the hidden states, see Methods. In this way, we are able to compute *P* (*σ*_*k*_ |*x*_1*:k*_) via the forward algorithm, and to sample backwards from *P* (*σ*_*k*_ |*x*_1*:T*_) with the forward-backward algorithm, for any *k* = 1, …, *T*. Using the formulation of the likelihood described in the Methods, and automatic differentiation to provide derivatives, we sample from the posterior distribution using an Hamiltonian Monte Carlo algorithm (***Neal, 2011***; ***Hoffman and Gelman, 2014***), implemented in Stan (***Carpenter et al., 2017***). On synthetic data simulated from the anaphase model of eq. (1), we are able to recover the true parameters used to simulate the data (Supp. Fig. 2), and infer the hidden microtubule attachment states and directional switches.

We demonstrate the automated annotation of hidden microtubule attachment states on a trajectory of a kinetochore pair in Figures 3A,B,C. Estimates for the probability of occupying each discrete hidden state at a given time point are shown in Fig. 3B, showing a clear transition to the anaphase state, *A*, around 230s (black line, Fig. 3B). Based on the sampled hidden states, intermediate states during directional switches can be identified, *i*.*e*. directional switches (between coherent states +− to −+, or −+ to +−), can occur via the intermediate incoherent states, −− or ++ (Fig. 3C). Where intermediate states cannot be identified (eg. +− followed by −+), we refer to the switch as a joint switch. We infer all the biophysical model parameters jointly; the inferred marginal distributions for each of the biophysical model parameters are shown in Fig. 3D. Based on trace plots and comparison between prior and posterior marginals, all model parameters satisfy practical identifiablity (***Hines et al., 2014***; ***Browning et al., 2020***), except *L* for which an informative prior was used, as (***Armond et al., 2015***), because of an identifiability issue, see Methods. The Bayesian framework allows us to propagate forward uncertainty in the parameter estimates, and thus simulate from the posterior predictive distribution, *p*(*x*_*t*_| *x*_1:(*t*−1)_, *σ*_*t*_), as shown in Fig. 3E. The true data points lie within the predictive interval from the model, indicating that the model describes the observed data.

**Figure 3.**
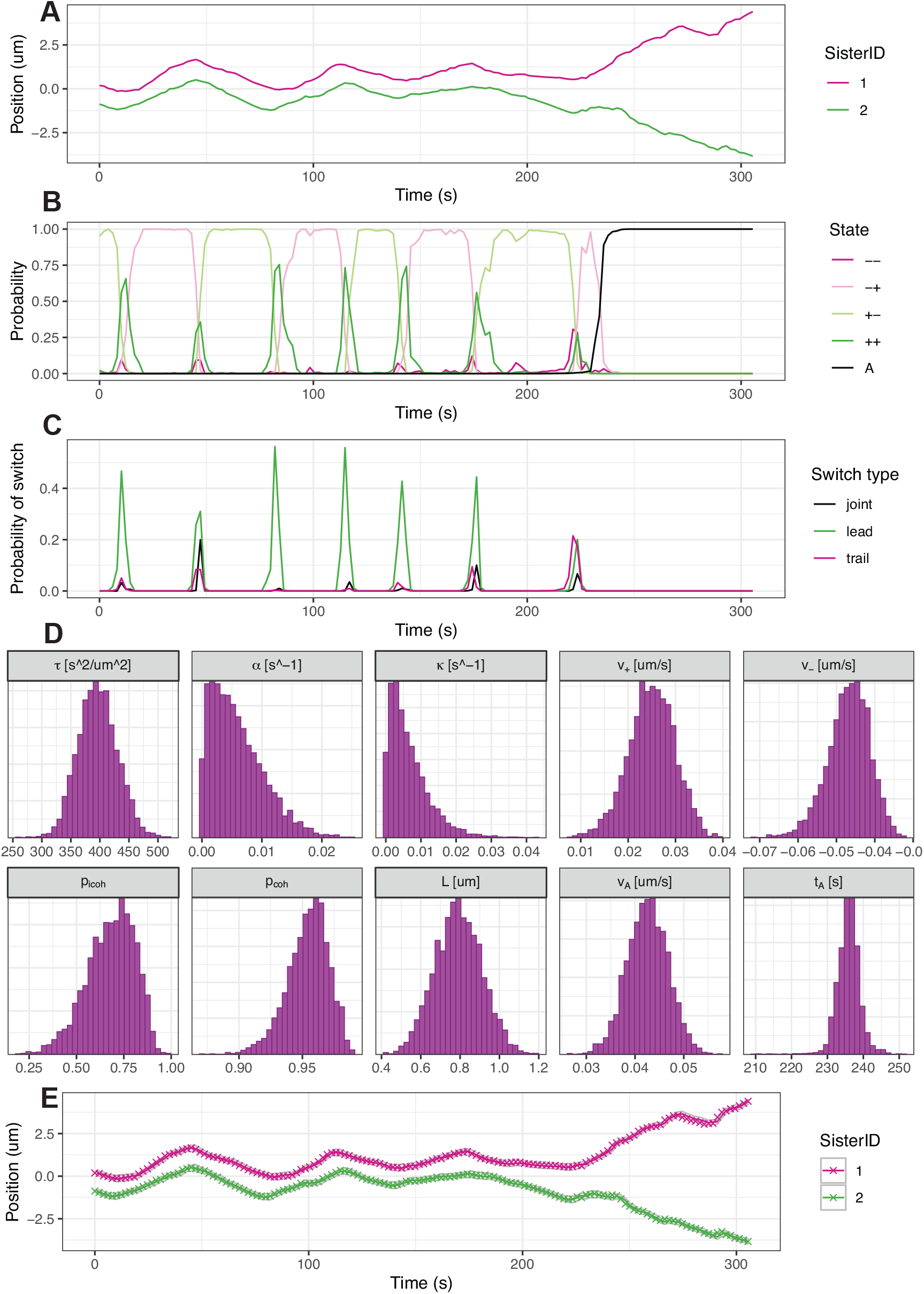
Bayesian inference enables automated annotation of microtubule attachment hidden states, and estimation of biophysical parameters (A) Observed track of kinetochore pair from experimental data. (B) Inferred probability of hidden states over time, *P* (*σ*_*t*_ |*x*_1*:T*_ ; *θ*) as sampled via the backward algorithm (see Methods). (C) Probability of a directional switch initiated by the leading kinetochore, trailing kinetochore, or a joint switch. Switching probability is assessed using the sampled hidden states and corresponds to a proportion of MCMC samples matching a particular pattern of states (eg. [−+,−+,++,+−] or equivalent for a LIDS) corresponding to a given switch type. D) Marginal posterior distribution of biophysical parameters for the trajectory data in (A). (E) Prediction from the filtering distribution *P* (*x*_*t*_ |*x*_1:(*t*−1)_, *σ*_*t*_). Coloured crosses indicate observed data.

### Incorporating anaphase directional reversals with a hierarchical model

One aspect of chromosome dynamics not captured by the model of the previous Section are transient reversals of the usual poleward motion during anaphase (see Figures 3E and 4C). These reversals have been observed in previous studies (***Skibbens et al., 1993***), whilst metaphase-like chromosome oscillations have been shown to persist into anaphase upon inhibition of protein dephosphorylation (***Su et al., 2016***). However, the cause of these reversals is not understood. Without reversals in the model of chromosome dynamics, there is a mismatch between the model and data due to misspecification of the model, resulting in MCMC convergence issues (Supp. Fig. 3) for some trajectories where reversals occur.

To incorporate reversals into the mechanical model of the previous section, we add an additional discrete hidden state, *R*, only accessible from the anaphase state, *A*, as shown in Fig. 4A. Since the forces acting on a kinetochore that cause, and act during, a reversal are not well understood, we model the reversal state dynamics as a pure diffusion:

**Figure 4.**
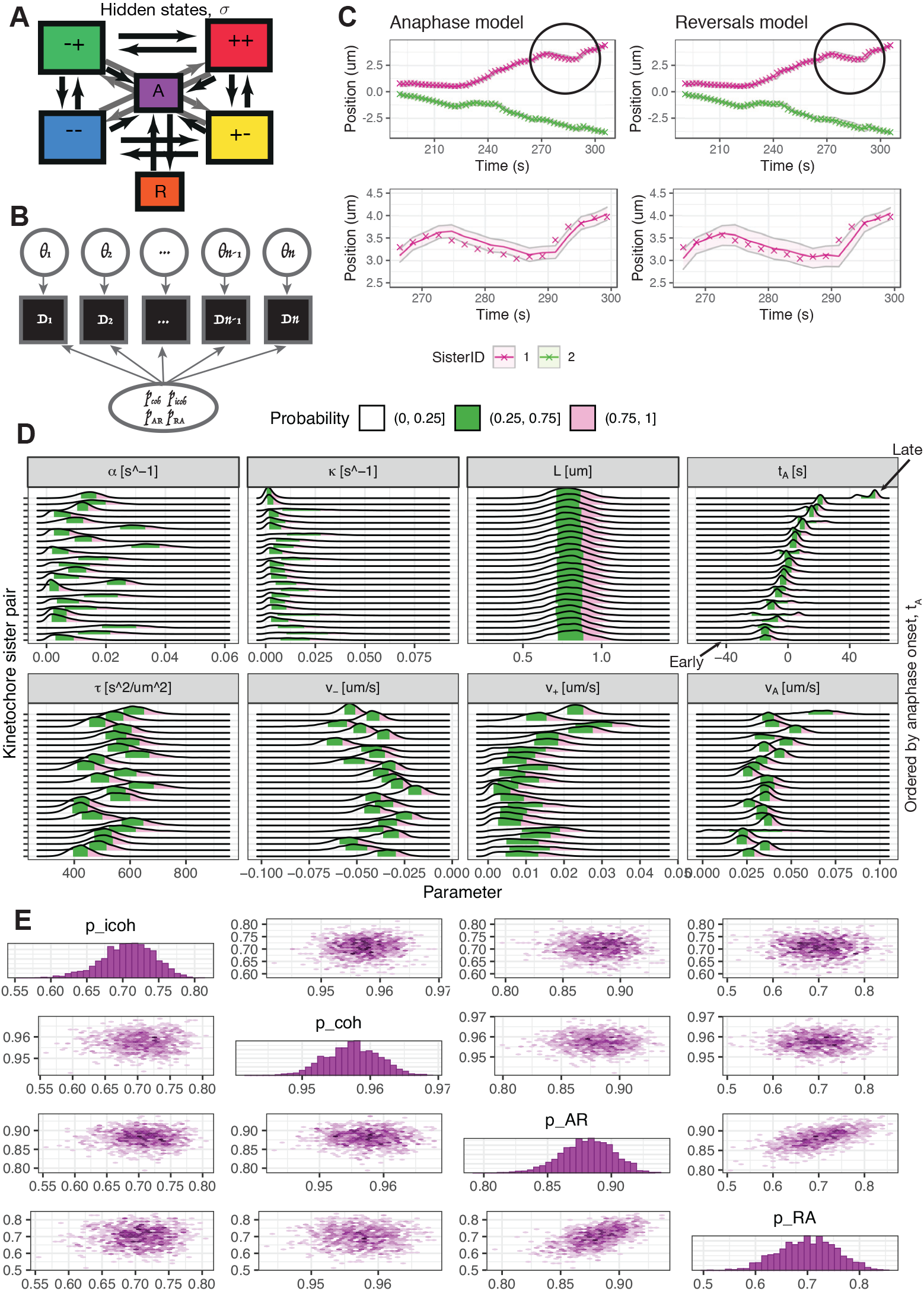
Hierarchical Bayesian framework for inference of rare reversals in anaphase. (A) Graphical structure of transitions between hidden states. All metaphase states, {++, +−, −+, −−}, are accessible from each other by either a single sister switching, or both sisters switching within a time step. The anaphase state, *A*, is accessible from all the metaphase states. From the anaphase state, *A*, transitions to and from the reversal state, *R*, are possible. (B) Schematic showing the hierarchical structure of shared rate parameters, *θ*_*SP*_ = (*p*_icoh_, *p*_coh_, *p*_*AR*_, *p*_*RA*_), and individual biophysical parameters, *θ*_*BP*_ = (*τ, α, κ, υ*_−_, *υ*_+_, *L*), unique to each kinetochore sister pair. (C) Prediction from the filtering distribution *P* (*x*_*t*_ |*x*_1:(*t*−1)_, *σ*_*t*_; *θ*) on a trajectory with reversals during anaphase. The reversal in anaphase is circled and the observed data (coloured crosses) lies outside the shaded region, the 95% credible region for the model predictions, between 270s and 290s for the simple biophysical anaphase model, but not for the hierarchical model. (D) Biophysical parameter marginal posterior distributions for individual kinetochore sister pairs. Pairs are ordered by the posterior mean time of anaphase onset, *t*_*A*_, which are shown relative to the median time of anaphase onset for the cell. The green shaded region highlights the interquartile range, while the grey and pink shaded regions show the lower and upper tails respectively. (E) Density plot of the pairwise posterior distribution for shared rate parameters, *p*_icoh_, *p*_coh_, *p*_*AR*_, *p*_*RA*_ for a single cell, showing correlations between parameters. Histograms of the marginal posterior distributions are shown on the diagonal.

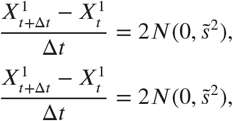

using a higher diffusion coefficient (we found the factor of 2 worked on our data). This ensures that a wide variety of reversal behaviours will be compatible with the model, and can be refined in future as mechanisms are identified. Crucially, the additional reversal state avoids model misspecification that would otherwise be present because of these unaccounted for reversal events, and thus avoids convergence issues of the MCMC algorithm.

However, a key complication is that reversals are relatively rare events and do not occur in every sister pair. In order to parametrise this model from experimental data, we need sufficient reversal events to infer the rate of transition from state *A* to state *R*, and vice versa. If fitted to data for a single pair with no reversals then there would be an identifiability problem. We therefore developed a hierarchical model for joint inference of all the (tracked) kinetochore pairs in a cell, with both shared parameters and individual kinetochore pair parameters. This enables estimation of rare event transition rates (shared parameters), including between states *A* and *R*, based on trajectories from all kinetochore sister pairs in the cell. A further advantage is that when pooling all the directional switching events for all kinetochore pairs in a cell, we have a large sample of switching events and thus obtain a tighter, more informative posterior distribution for the switching parameters compared to inference on individual sister pair trajectories.

The graphical structure of the hierarchical model with shared transition rates is shown in Fig. 4B. The transition probabilities between the hidden states are <u>s</u>hared <u>p</u>arameters relevant to all kinetochore pairs in a cell, *θ*_*SP*_ = (*p*_icoh_, *p*_coh_, *p*_*AR*_, *p*_*RA*_), where *p*_*AR*_ is the probability of remaining^1^ in the anaphase state, *A*, over a time step Δ*t* (and similarly for *p*_*RA*_ the probability of remaining in the reversal state). The remaining <u>b</u>iophysical <u>p</u>arameters *θ*_*BP*_ = (*τ, κ, α, L, υ*_−_, *υ*_+_, *υ*_*A*_, *t*_*A*_) are assumed to be unique to each kinetochore pair and are inferred independently for each pair. Several of these biophysical parameters have been shown to vary based on position in the metaphase plate (***Armond et al., 2015***) which motivates keeping independent parameters for each sister kinetochore pair.

We demonstrate inference for the hierarchical model based on data from all tracked pairs in a single cell (Fig. 4C-E); inferred hidden states including the reversal state, *R* are shown for some representative trajectories in Supp. Fig. 4. The marginal posterior distribution of the individual biophysical model parameters for each kinetochore sister pair are shown in Fig. 4D, with the posterior distribution of the shared rate parameters shown in Fig. 4E to visualise marginal distributions and pairwise relationships between rate parameters. We obtain tight estimates of all parameters; recall that the natural spring length parameter, *L*, has a strong informative prior as described in Methods. In particular the switching rates for metaphase oscillations are tighter than for single trajectory pairs (posterior standard deviation 4.1 and 3.5 times smaller for *p*_coh_ and *p*_icoh_ respectively for the trajectory shown in Fig. 3A), reflecting the greater number of directional switching events in a cell, whilst the reversal transition rate *p*_*AR*_ is also well inferred. The transition from *A* to *R* is rare hence the probability of remaining in the *A* state per frame, *p*_*AR*_ is close to 1. This model is consistent with observations with reversals being rare events; only 6% of kinetochore pairs spend on average more than 10 frames in the reversal state (Supp. Fig. 5).

The hierarchical model’s predictions are better than those from the model without reversals. Specifically, we compared the posterior predictive distribution, *p*(*x*_*t*_ | *x*_1:(*t*−1)_, *σ*_*t*_) for the cell based hierarchical model with shared rate parameters and the previous kinetochore pair based metaphaseanaphase model. For a trajectory with a reversal, we find the observed data (coloured crosses) lies within the credible region for the hierarchical model (Fig. 4C), whereas the data lies outside the model’s credible region for the simpler anaphase model during the reversal (Fig. 3E, and Fig. 4C). We observe heterogeneity between different kinetochore sister pairs in cell (Fig. 4D). Ordering the kinetochore sister pairs by the relative time of anaphase onset, *t*_*A*_, reveals correlations with parameters such as *τ* and *υ*_*A*_ (Fig. 4D, Supp. Fig. 6). The velocity in anaphase, *υ*_*A*_ and the precision parameter, *τ*, both show an increase among pairs with later anaphase onset times; increased velocity in anaphase for late separating pairs is consistent with previous studies (***Armond et al., 2019***).

### Distribution of biophysical parameters across the population of kinetochores

To assess heterogeneity of kinetochore dynamics in a population of non-transformed human RPE1 cells, we consider the distribution of posterior median parameter estimates for the population of kinetochore pairs tracked in *N* = 25 cells. This distribution is shown in Fig. 5 for both the individual parameters unique to each kinetochore pair (Fig. 5A, summarised in Table 1), and the rate parameters shared across cells (Fig. 5B). We exclude cells and kinetochore pairs for which diagnostics of the MCMC chains indicate that convergence has failed (see Methods and Supp. Fig. 7).

**Table 1.**
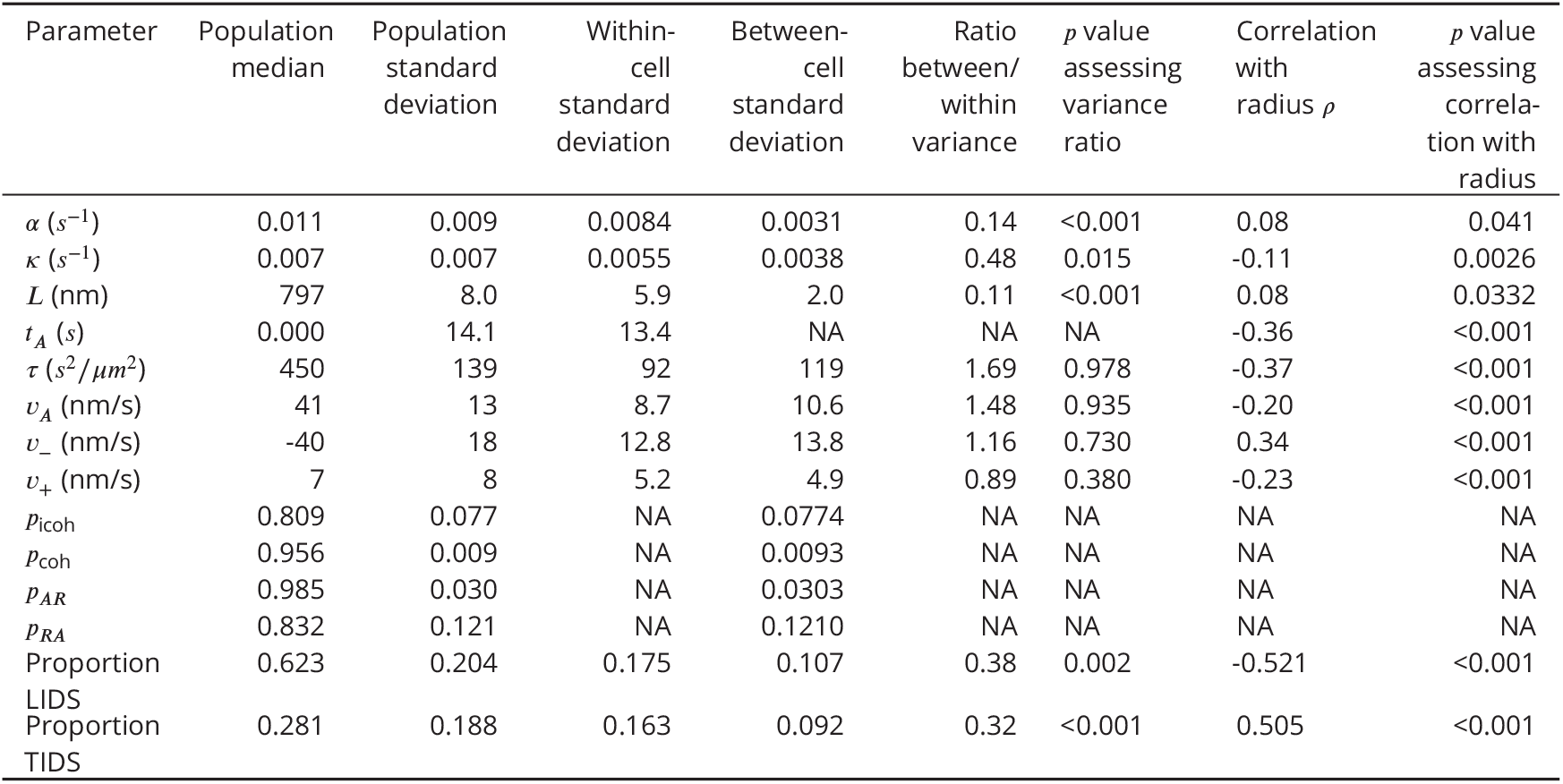
Estimates of biophysical parameters in RPE1 cells, and correlation with radial position in metaphase plate based on *n* = 684 kinetochore pairs from *N* = 25 cells

**Figure 5.**
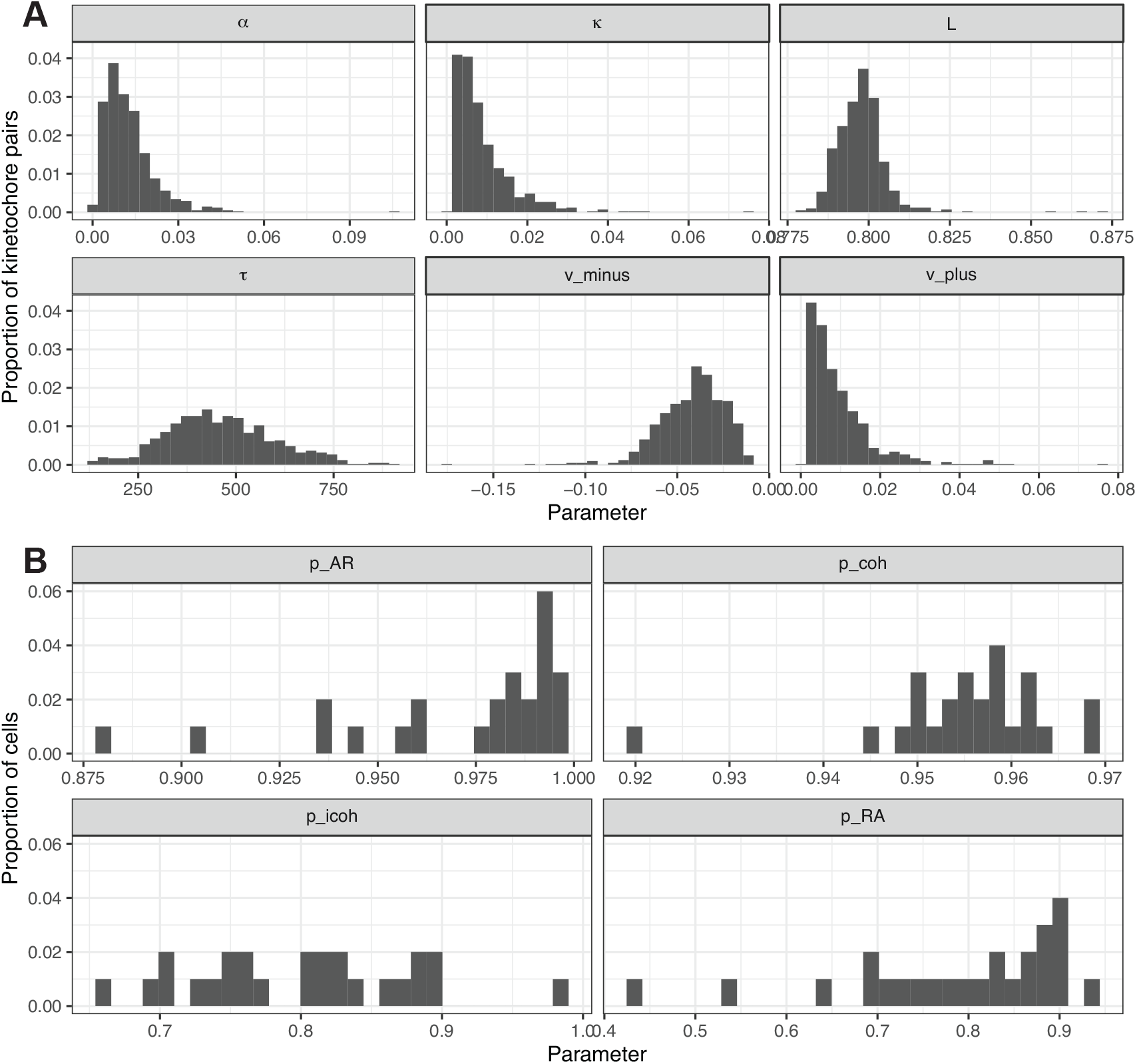
Distribution of model parameters for a population of *n* = 684 kinetochore pairs from *N* = 25 cells. (A) Histograms of the median posterior estimate for biophysical parameters *α, κ, L, τ, υ*_−_, *υ*_+_ across a population of trajectories from multiple cells. (B) Histograms of the median posterior estimate for each of the shared rate parameters, *θ*_*SP*_ = (*p*_icoh_, *p*_coh_, *p*_*AR*_, *p*_*RA*_), across the population of cells.

The main difference to highlight compared to previous estimates based on HeLa cells (***Armond et al., 2015***) is the distribution of the spring constant, *κ*, which has shifted to smaller values for RPE1 cells (0.009 ± 0.007 *s*^−1^; mean ± population standard deviation) compared to the previous estimates for HeLa cells (0.03 ± 0.01 *s*^−1^). This therefore indicates that RPE1 cells have a more compliant (weaker) centromeric chromatin spring than for HeLa cells. Other notable differences compared to previous estimates in HeLa cells relate to the microtubule speed parameters, *υ*_−_ and *υ*_+_. The magnitude of these parameters have a greater difference (| *υ*_−_| −*υ*_+_) in RPE1 cells compared to HeLa cells; *υ*_−_ is estimated as −40 ± 18 nm/s in RPE1 cells versus −35 ± 15 nm/s in HeLa, similarly *υ*_+_ is estimated as 7±8 nm/s in RPE1 cells versus 13± 16 nm/s in HeLa, where mean values ± standard deviation of the population are given. It should be noted however that the HeLa cell analysis in ***Armond et al. (2015***) is based on far lower coverage of kinetochores than the current analysis.

To assess whether there are any links between biophysical parameters and segregation errors, we considered laziness of kinetochores, as defined in ***Sen et al. (2021***). Lazy kinetochores show impaired segregation compared to other kinetochores that segregate to the same daughter cell. We found that kinetochores with high laziness have significantly higher values for *υ*_+_ (*p* = 0.02, see Supp. Fig. 8) compared to the remaining population of kinetochores. No other biophysical variables were significant.

The parameter distributions over the cell population demonstrate heterogeneity between cells, Fig. 5. Since heterogeneity is observed within cells, for instance, trends with metaphase plate location (see Table 1 and Supp. Fig. 9), it is unclear how much of population heterogeneity derives from variation within cells. To determine whether variation between cells or within cells contributes more to the variation, for each parameter we consider between-cell variation: the standard deviation across the population of the median parameter estimates per cell (median over kinetochore pairs in a cell); versus samples of the within-cell variation: the parameter standard deviation of each cell’s kinetochore pairs (Fig. 6A). As indicated in Fig. 6B and Table 1, we find greater within-cell variation compared to between-cell variation, particularly for the polar ejection force parameter, *α*, and the spring parameters, *L* and *κ*. This is evident because the cell-to-cell variation (red dot) lies below the within-cell variability (black dots) for these parameters. Assessing the ratio of between-cell variance to within-cell variance and whether this differs from 1 with an *F* -test indicates a significant difference for the polar ejection force parameter, *α*, (*p* < 0.001), the natural spring length, *L*, (*p* < 0.001), and spring constant, *κ*, (*p* = 0.015). Although this result may be expected for the natural spring length, *L*, which is subject to a strong informative prior distribution and thus has near zero variation between cells, it is perhaps surprising that the PEF parameter, *α*, and spring constant, *κ*, have a prescribed variation over the metaplate that is similar in different cells.

**Figure 6.**
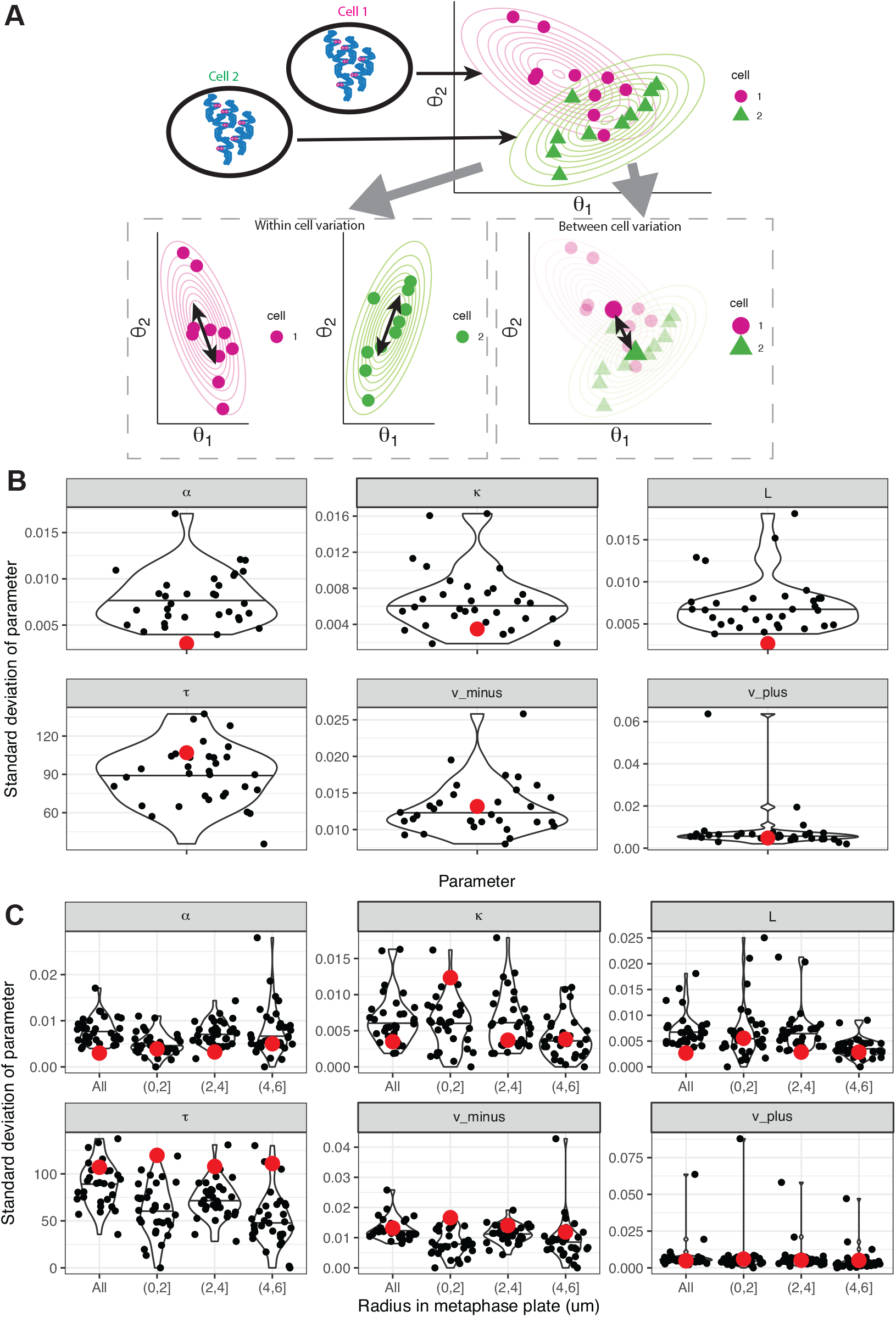
Comparison of heterogeneity in biophysical parameters between and within cells. (A) Schematic diagram defining within cell variation and between cell variation. For two cells (green and magenta), their estimated parameter values for *θ*_1_ and *θ*_2_ are shown by magenta circles and green triangles respectively. The standard deviation of *θ*_*i*_ for the magenta and green points respectively quantifies the variation within a cell (left), whereas taking a cell average and considering the standard deviation between cells quantifies the variation between cells. (B) Violin plots of the standard deviation of the variation between kinetochore pairs within each cell for biophysical parameters *α, κ, L, τ, υ*_−_, *υ*_+_. Each black dot corresponds to the variation within a cell. Marked red dots are the standard deviation of the median parameter estimates for each cell showing the level of between-cell heterogeneity. (C) Violin plots as in (B) showing the full data (all kinetochores) and subpopulations of kinetochores in the indicated ranges for the radial position in the metaphase plate. Assuming a normal distribution for within-cell variation and evaluating the percentile of the between-cell variation, for *α* this gives *p* = 0.044 for all kinetochores, but is no longer significant for subpopulations of kinetochores at particular radial locations. Results based on *n* = 684 kinetochore pairs from *N* = 25 cells.

Since spatial variation of the PEF was previously demonstrated by ***Armond et al. (2015***), we therefore examined the extent to which this heterogeneity can be explained by subpopulations of kinetochores located in particular spatial locations of the metaphase plate. We compared the same heterogeneity statistics for kinetochores located, on average, 0 to 2 *µ*m, 2 to 4 *µ*m, and 4 to 6 *µ*m from the centre of the metaphase plate (Fig. 6C). For parameter *α*, between-cell variation (red dot) was below the within-cell variability (black dots) when considering all kinetochore pairs, but on the radial kinetochore subsets the red dot lies within the cell variation distribution (black dots) suggesting that heterogeneity within this population and between cells are similar for these subsets. In contrast, after accounting for radius, the red dot lies above the black points for the precision parameter, *τ*, suggesting that, once radius is accounted for, the noise varies much more between cells than within cells. These results indicate that radial position in the metaphase plate dictates kinetochore dynamic behaviour, making a notable contribution to variability in biophysical properties.

### Directional switching events vary spatially across the metaphase plate

Given the spatial variation in the biophysical parameters (see Table 1), we reasoned that there may also be effects on directional switching of chromosomes. Metaphase quasi-periodic oscillations require that both sister kinetochores change direction at a directional switch. A key distinction is between which sister switches first specifically switching events initiated by the leading kinetochore sister (lead induced directional switch, or LIDS, events), and those initiated by the trailing kinetochore sister (trail induced directional switch, or TIDS, events) are observed (***Armond et al., 2015***; ***Burroughs et al., 2015***). From the annotation of kinetochore trajectories, we can identify directional switching events and determine which sister initiated that switch, and subsequently correlate LIDS/TIDS events with our trajectory summary statistics, Fig. 7.

**Figure 7.**
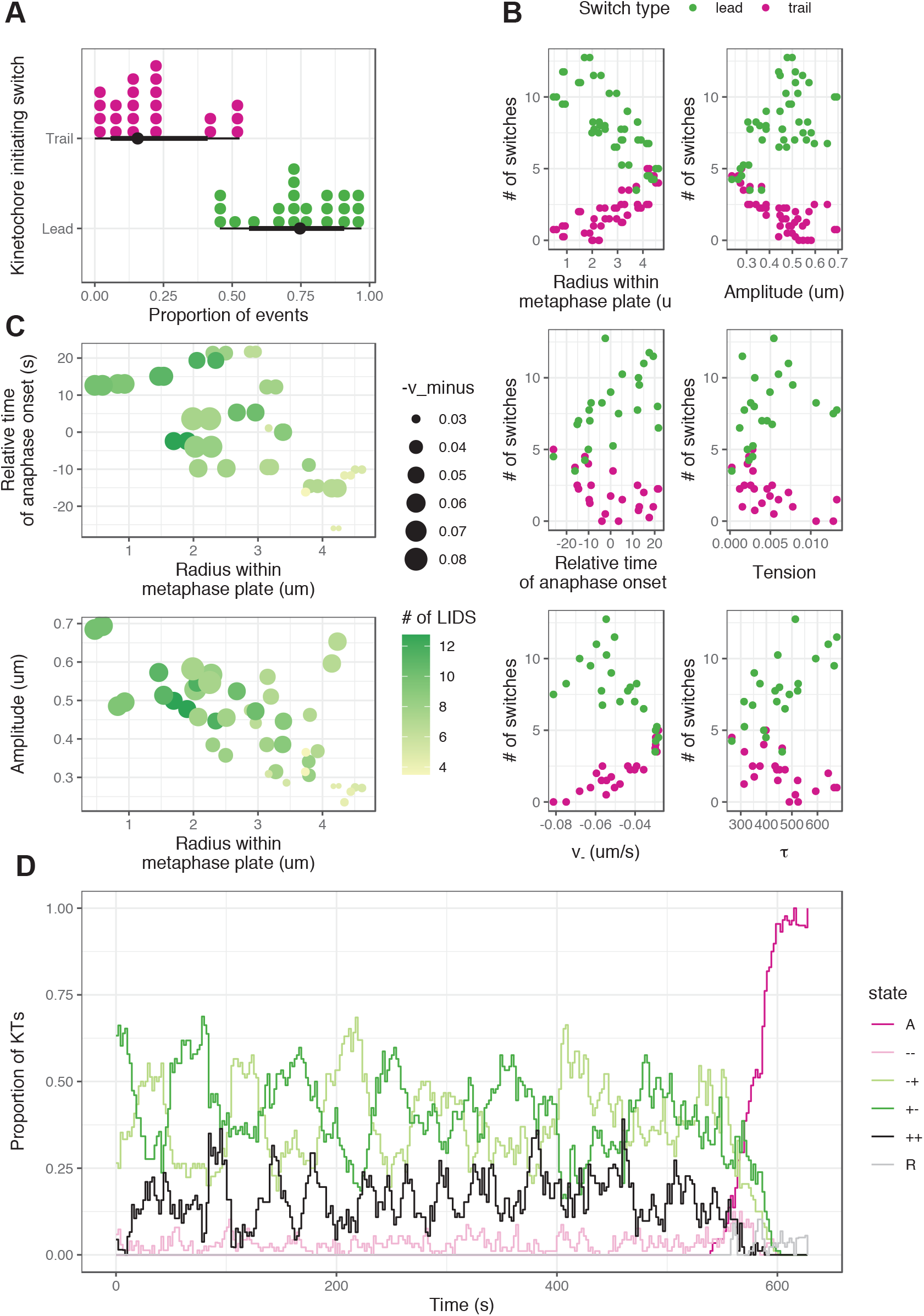
Directional switches of oscillating chromosomes vary across the metaphase plate. (A) Fraction of LIDS (green) and TIDS (pink) events as a proportion of the total number of switching events including joint switches. Each kinetochore pair gives rise to a LIDS and TIDS dot. (B) Relationship between the number of directional switches initiated by the leading (green) or trailing (pink) kinetochore sister, and other summary statistics describing the oscillatory dynamics. (C) Relationship between the number of LIDS events and other summary statistics indicating that many of these variables change together based on spatial position of kinetochore pairs within the metaphase plate. (D) Proportion of kinetochore sister pairs in a given hidden state at each time point. All data shown in this figure relate to a single cell; another cell is shown in Supp. Fig. 10 and population summaries are given in Table 1.

By fitting the hierarchical model of metaphase-anaphase dynamics to data from *N* = 25 cells at a time resolution of Δ*t* = 2.05s per image stack, and using forward filtering backward sampling (as described in Methods), we are able to sample from the distribution of hidden states through a trajectory. Given a sequence of states, a directional switch is defined as a pattern of states that changes between consecutive coherent runs moving in the opposite directions (+− to −+ or vice versa), which can have intermediate incoherent states ++ or −−, the sister kinetochores then moving in opposite directions. For example the sequence [+−, +−, +−, +−, −−, −+, −+, −+, −+] would be assessed as containing a TIDS, as would a sequence with a longer incoherent sequence such as [+−, +−, +−, +−, −−, −−, −−, −+, −+, −+, −+]. We averaged over MCMC samples and obtained the posterior number of LIDS and TIDS events in a sister kinetochore pair trajectory. This allows the proportion of LIDS and TIDS events to be computed, relative to the total number of events (Fig. 7A, Table 1).

Switch events in RPE1 cells are approximately two times more likely to be initiated by the leading kinetochore sister than the trailing kinetochore sister (Fig. 7A), consistent with previous results in HeLa cells (***Armond et al., 2015***; ***Burroughs et al., 2015***). Furthermore, we find that kinetochore pairs close to the centre of the metaphase plate have more LIDS events and fewer TIDS events, as shown in Fig. 7B and Table 1. Many other summary statistics and model parameters including oscillation amplitude, the relative time of chromatid separation, tension, microtubule speed in the (leading) depolymerization state *υ*_−_, and the noise parameter, *τ*, all show strong correlations with the number of LIDS events (Fig. 7B). However, many of these covariates vary together spatially within the metaphase plate (Fig. 7C) so determining causality of the factors responsible for these effects on switching dynamics is unclear. Moreover, as for the biophysical parameters in the previous section, we observe significantly higher variation in the proportion of LIDS and TIDS events within cells than between cells (Table 1) (*F* -test; LIDS: *p* = 0.002, TIDS: *p* < 0.001).

When considering the proportion of kinetochore pairs in a given state over time (averaged over a cell’s kinetochore population), we find oscillatory behaviour (Fig. 7D). These oscillations occur on a similar timescale to the oscillations for single kinetochore pairs, perhaps suggesting coordination across the population in the kinetochore oscillations, similar to the correlation between non-sister kinetochore trajectories observed in ***Vladimirou et al. (2013***). Intuitively we would expect that if kinetochore pair oscillations were independent, the proportion of kinetochore pairs in each state would be constant over time. Simulations from a 4 state Markov model with spontaneous switching between states (see Methods) also exhibit fluctuations in the proportion of kinetochore pairs in each state (Supp. Fig. 11), although lack the regular periodicity observed in the experimental data. Thus, the oscillations seen in the average state proportion, Fig. 7D, are likely a result of averaging over a finite number of oscillating kinetochore pairs (46 in human cells, and 46 in Supp. Fig. 11); in the limit of infinite kinetochore pairs these fluctuations would disappear.

Finally, examination of the K-fibre polymerisation hidden states around anaphase in Fig. 7D shows that no particular state dominates - thus anaphase onset is not coordinated with a particular metaphase oscillation phase. There is only a small increased proportion of the (−−) states in the lead up to anaphase (Fig. 7D, light pink line). Whether this is mechanistically relevant is unclear, as (−−) is similar to the anaphase state in that both sisters are depolymerising.

### Coordination of directional switching events in space and time

To further assess the extent of the coordination of directional switching across the metaphase plate, we considered the influence of directional switching events on the switching of neighbouring kinetochore pairs. Taking a dataset of directional switching events in a cell, as determined by the metaphase-anaphase model, we use a self-exciting point process model known as a Hawkes process (***Hawkes, 1971***; ***Reinhart, 2018***) (Fig. 8A) to determine how contagious directional switching events are. Label the switch events by *i*, then a switch event is given by (*j*_*i*_, *t*_*i*_, ***s***_*i*_), the event occurring in pair *j*_*i*_ at time *t*_*i*_, kinetochore position ***s***_*i*_. The Hawkes process intensity for a switch event at time *t*, position ***s***, kinetochore sister pair *j* is conditioned on the history of all previous switches, *t*_*i*_ < *t, i* = 1, 2, and given by

**Figure 8.**
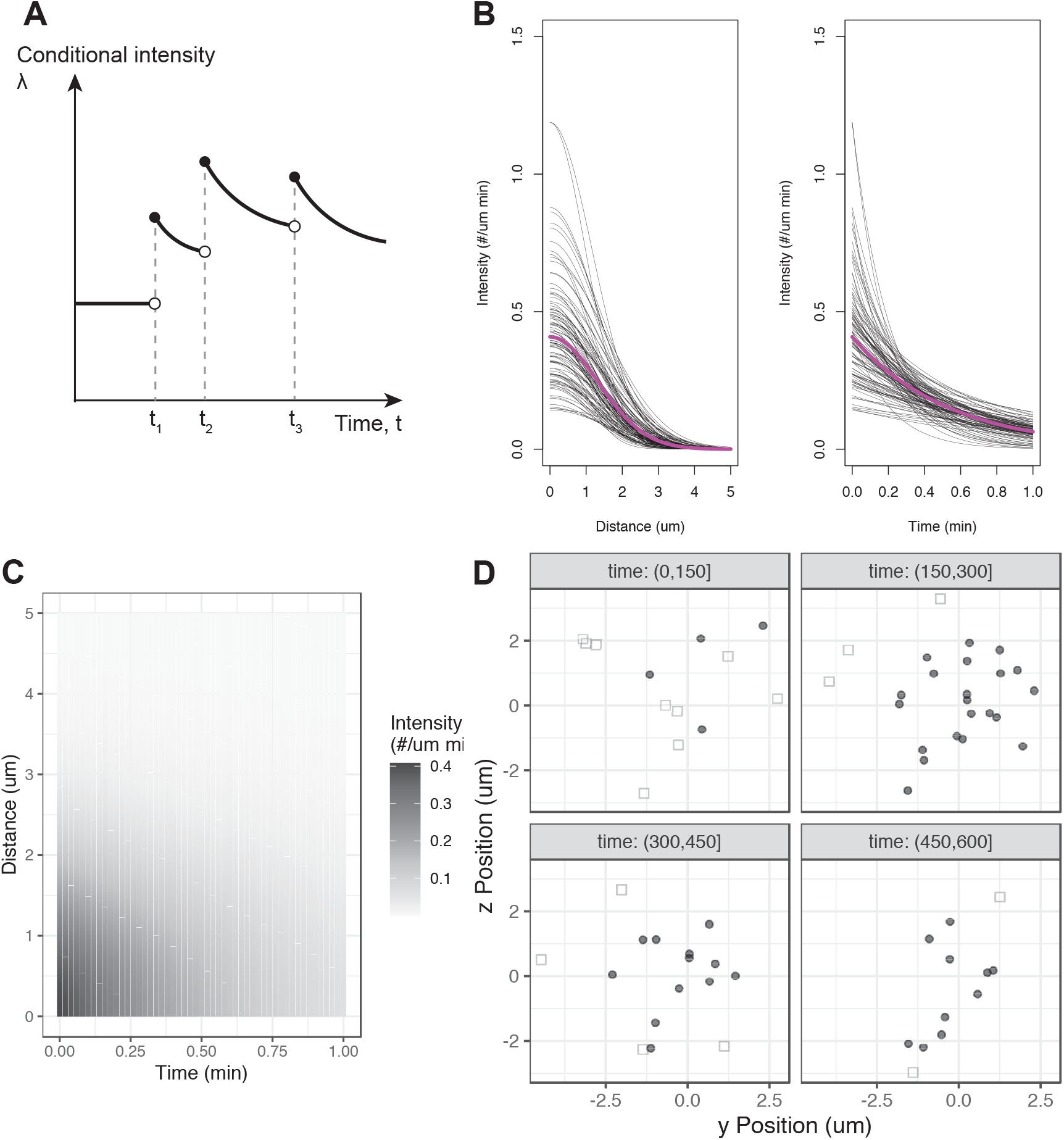
Directional switching events are coordinated in space and time, as revealed by a Hawkes process model. (A) Schematic of conditional intensity with a Hawkes process model (assuming only self-excitory behaviour in time, and no spatial kernel) showing increase in the event rate after an event, the effect decaying with time from the event. (B) The self-excitatory component of the Hawkes process showing decay in interactions between switching events in space and time. The spatial kernel is visualised at *t* = 0, while the temporal kernel is visualised at a distance of 0. Each black line corresponds to a sample from the posterior, while the magenta line corresponds to the posterior mean. (C) Self-excitatory component of the Hawkes process visualised in space and time. (D) Example cell with switching events classified as excitatory (filled circles) or as background (open squares). The *y* and *z* positions shown correspond to positions within the metaphase plate. Subplots divide metaphase into sections of 150s. Data relates to a single cell.

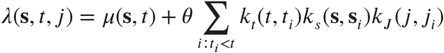

where *k*_*t*_(*t, t*_*i*_) is the (exponential) kernel in time, *k*_*s*_(***s, s***_*i*_) is the (Gaussian) kernel in space, and *k*_*J*_ (*j, j*_*i*_) is a kernel that removes interactions between sister kinetochores. We use an exponential decay kernel for influence of events in the past, and a Gaussian kernel to discount for the 3D distance of that event - the strengths of both of these kernels are inferred from the data. A point process without self-exciting interactions would have only the *µ*(***s***, *t*) term, while the sum term represents the influence of previous switching events.

Fitting the Hawkes process in a Bayesian framework (details in Methods) allows us to assess whether one directional switch is likely to be coordinated with other directional switches locally in space and time. This analysis revealed interactions between switching events over a timescale of 36s [18.9,84.1] and a lengthscale of 1.3*µ*m [1.02,1.66], where square brackets indicate the 95% credible interval, as shown in Fig. 8B and C. The parameter *θ* represents the average number of switching events triggered by a single switching event, and this was estimated as 0.72 [0.43,1.05]; since this is less than 1 the event structure in space and time comprises small clusters that are excited from a spontaneous event, the cluster size being stochastic. From 100 switching events, we would thus expect 72 switching events to be triggered by these, which will subsequently trigger further switches. To visualise this, we classify events triggered by other switching events (and thus a result of excitatory behaviour) as those with the highest estimated conditional intensity. This allows us, in Fig. 8D, to highlight the switching events influenced by other switching events by excitatory behaviour (filled circles) versus switching events that occur spontaneously due to the background rate of switching (open squares). Thus, directional switching events of neighbouring kinetochore pairs are spatio-temporally coordinated.

### Spatial coordination of K-fibre polymerisation and depolymerisation states

Although the Hawkes process demonstrates an influence of switching events of other kinetochore pairs on the switching rate of an individual kinetochore, it does not allow for the nature of the switch or directionality of the kinetochores. To further investigate local coordination of kinetochore state, and related movements, we define a measure of alignment between the hidden states of a kinetochore pair (polymerising/depolymerising for each K-fibre) and that of neighbouring kinetochore pairs. Specifically, the alignment between kinetochore pair *i* and pair *j* at time *t* is defined as

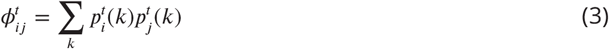

where 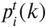 is the probability that pair *i* is in state *k* at time *t*. Based on average positions of kinetochore pairs within the metaphase plate, we identify *k* nearest neighbours of a kinetochore pair (Fig. 9A), and average the alignment of neighbouring kinetochore pairs as follows

**Figure 9.**
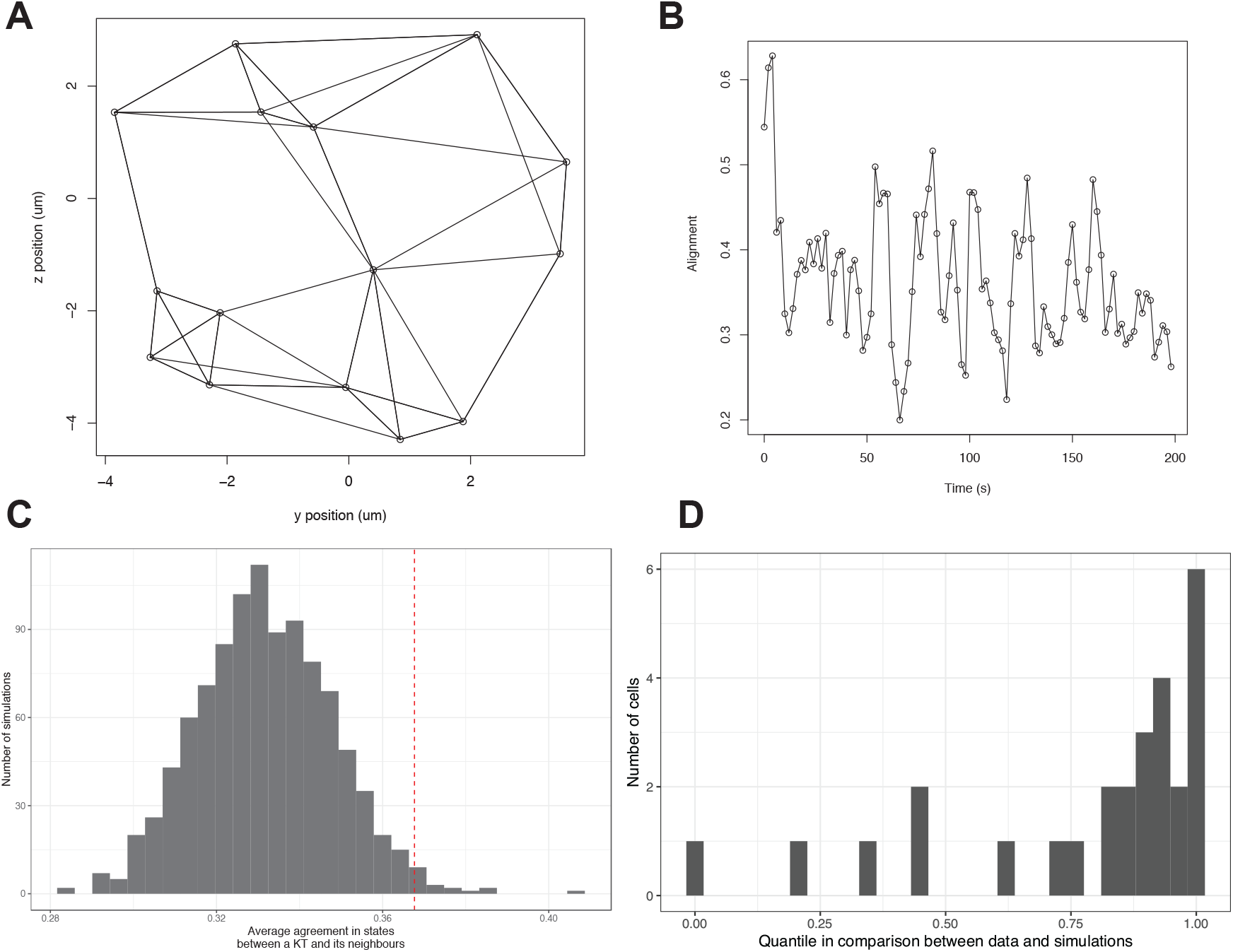
Local coordination of K-fibre polymerisation and depolymerisation states. (A) Network of *k* = 4 nearest neighbours of kinetochore pairs based on average positions within the metaphase plate. (B) Average alignment of states, *ϕ*^*t*^, between neighbouring kinetochore pairs oscillates over time. (C) Comparison of average alignment from experimental data with simulations from a 4 state Markov process. Parameters for the Markov process are *p*_coh_ = 0.96, *p*_icoh_ = 0.83 are obtained from the posterior median estimate from fitting to this cell. The grey histogram shows average alignment computed for 1000 model simulations, while the dashed red line shows average alignment evaluated for experimental data. (D) Histogram of the percentile obtained from comparison between simulation and observed data for the average alignment for each cell in a population of *N* = 25 cells. A large proportion of percentiles are close to 1 indicating higher agreement of states in the observed data than expected from simulations of a null model. Plots (A-C) refer to the same example cell; data in (D) is based on *N* = 25 cells.

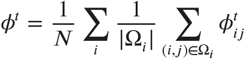

where Ω_*i*_ is the set of *k* nearest neighbours for a kinetochore pair *i* and *N* is the total number of pairs. Full alignment between all kinetochore pairs would give *ϕ*^*t*^ = 1. Using this measure of alignment, we find that neighbouring kinetochore pairs have greater average state alignment in the data than can be accounted for based on simulations from a model without any interactions (Fig. 9C,D). This suggests that interactions between kinetochore pairs are required to account for the alignment that we observe.

The alignment, *ϕ*^*t*^, of all pairs in a cell oscillates over time, as shown in Fig. 9B. Taking the autocorrelation in time of the average alignment of a kinetochore with its neighbours shows that these oscillations have a frequency similar to that of kinetochore pair breathing (≈ 20s). This indicates that neighbouring kinetochores interact, influencing each other so that there is local coordination of direction of movement, potentially via interactions between their respective K-fibres (***Tolić, 2018***; ***Simunić and Tolić, 2016***).

### Spatial coordination of anaphase onset

To determine if this local coordination of metaphase dynamics extends to local coordination of chromosome segregation, we analysed whether there is evidence of spatial coordination in anaphase onset. Taking the anaphase onset times inferred from the hierarchical metaphase-anaphase model, Fig. 4, we ordered the kinetochore pairs by increasing anaphase onset time. With this ordering, we visualised the spatial positions of kinetochore pairs over the sequence of anaphase onset times in a cell (Fig. 10A) and observed that kinetochore pairs close together in space tend to enter anaphase close together in time. To quantify this, we computed the 2D Euclidean distance within the metaphase plate between kinetochore pairs, and the difference between their separation times, *t*_*A*_ (Fig. 10B). We assessed whether there are a disproportionate number of interactions between kinetochore pairs that are close in space and time, (within 2um in space and 2s, or one frame, in time), compared to what would be expected from randomly permuting the times of anaphase onset of pairs, (Fig. 10C). This is quantified via the empirical cumulative distribution function (eCDF), indicating the percentile at which the experimental data lies in comparison with random permutations. Based on a population of cells, a much larger proportion of cells have a percentile close to 1 than would be expected (*p* = 0.007, KS test comparing to uniform distribution) as shown in Fig. 10D. These results indicate that there are local interactions between kinetochore pairs at anaphase onset, or that several pairs in similar spatial locations are all influenced by the same external factors (eg. separase activity).

**Figure 10.**
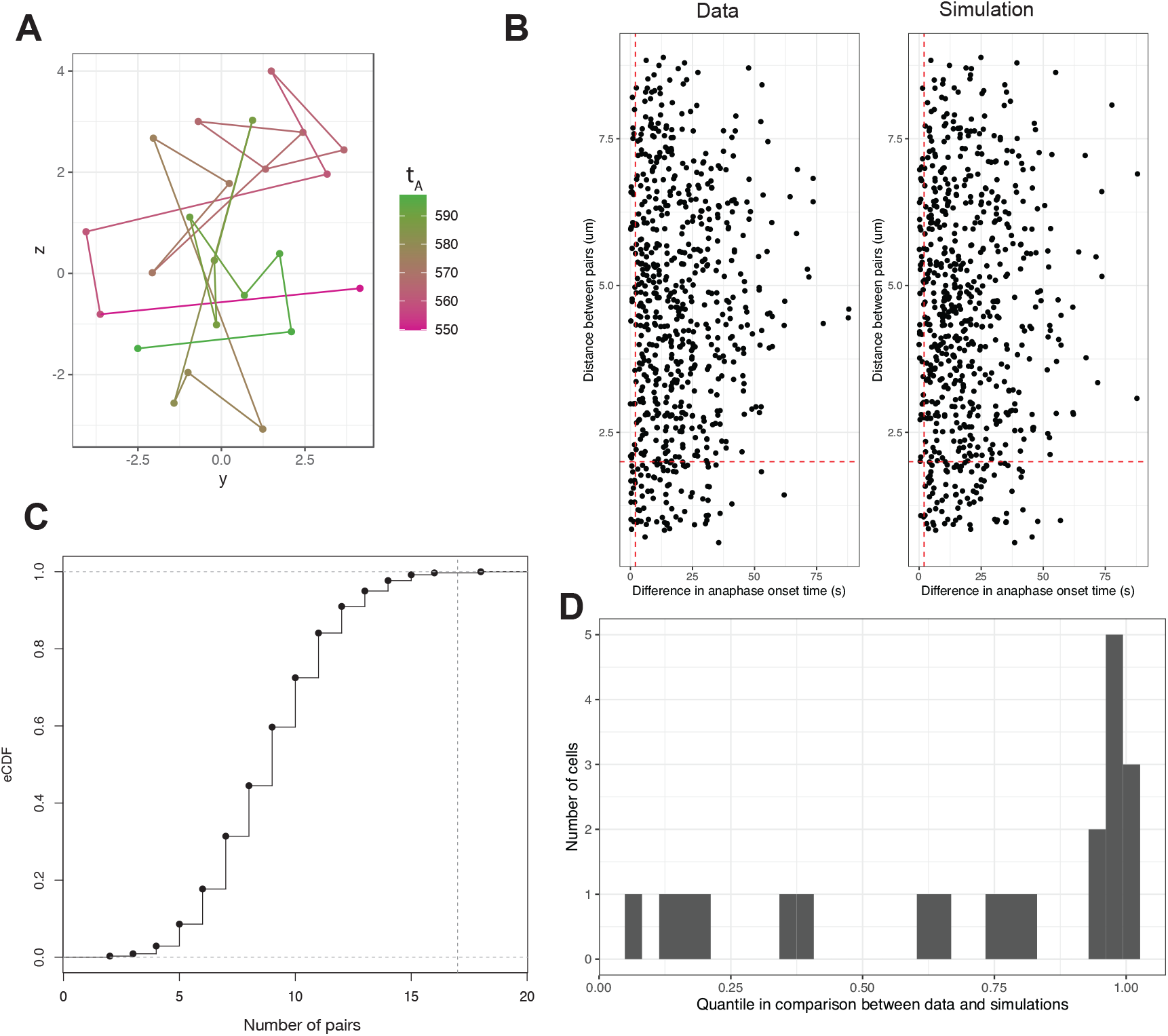
Anaphase onset times of kinetochore pairs exhibit local coordination. (A) Metaphase plate positions of kinetochore pairs in a cell at median anaphase onset time and coloured by time of anaphase onset *t*_*A*_ for each pair. Successive pairs in the anaphase onset time ordering are connected by lines. (B) Scatterplot of distances between all kinetochore pairs plotted against differences in their time of anaphase onset, *t*_*A*_, in observed data (left) and simulated data (right) via randomly permuting the observed distances. Red dashed lines indicate a region of interest at the left bottom corner of the plot representing pairs close in space and time (within 2s and 2um). (C) Empirical cumulative density function (eCDF) of number of pairs close in both space and time (bottom left corner in (B)) in random permutations of the times of anaphase onset. Vertical dashed line indicates where the observed data lies in comparison. (D) Histogram of (empirical) percentiles (as in (C)) associated with comparison between the number of pairs close in both space and time (bottom left corner in (B)) in observed data versus simulations (obtained by randomly permuting times of anaphase onset for all pairs in a cell). Plots shown in (A), (B) and (C) are from a single cell, while (D) shows data from *N* = 25 cells.

### Kinetochore pairs at the edge of the metaphase plate enter anaphase earlier than pairs at the centre

The onset of anaphase, when kinetochore pairs begin to separate and segregate towards their respective spindle poles, is tightly controlled temporally (***Holt et al., 2008***) and appears entirely synchronous at low time resolution. An asynchrony in anaphase onset times has previously been observed with a difference between first and last separation times of 60-90s in RPE1 cells (***Armond et al., 2019***; ***Sen et al., 2021***). Using our high time resolution imaging and inferred anaphase onset times *t*_*A*_, we confirm that kinetochore pairs in different parts of the metaphase plate separate at different times, with pairs at the edge of the plate separating on average over 20s earlier than pairs at the centre (Fig. 11A).

Furthermore, estimated anaphase speeds, *υ*_*A*_, are lower for kinetochore pairs at the edge of the plate (Fig. 11B). However, the same relationship is not seen for the framewise anaphase speed in 3D (Fig. 11C). By considering (a simple approximation of) the geometry of the mitotic spindle (see Fig. 11D), we can account for this. We assume that the spindle at anaphase onset can be approximated by two cones joined at their flat faces; these faces being the metaphase plate. This assumption ignores the curvature and chirality of the spindle (***Novak et al., 2018***; ***Pavin and Tolić, 2020***), but captures the key part of the spindle geometry relevant here. Consider the triangle between a K-fibre to a kinetochore at the centre of the metaphase plate, and the K-fibre to a kinetochore at the edge of the plate. The K-fibre at the edge is a distance *R* from the centre, and the K-fibre at the centre is a distance *S* from the pole, equal to the half spindle length.

**Figure 11.**
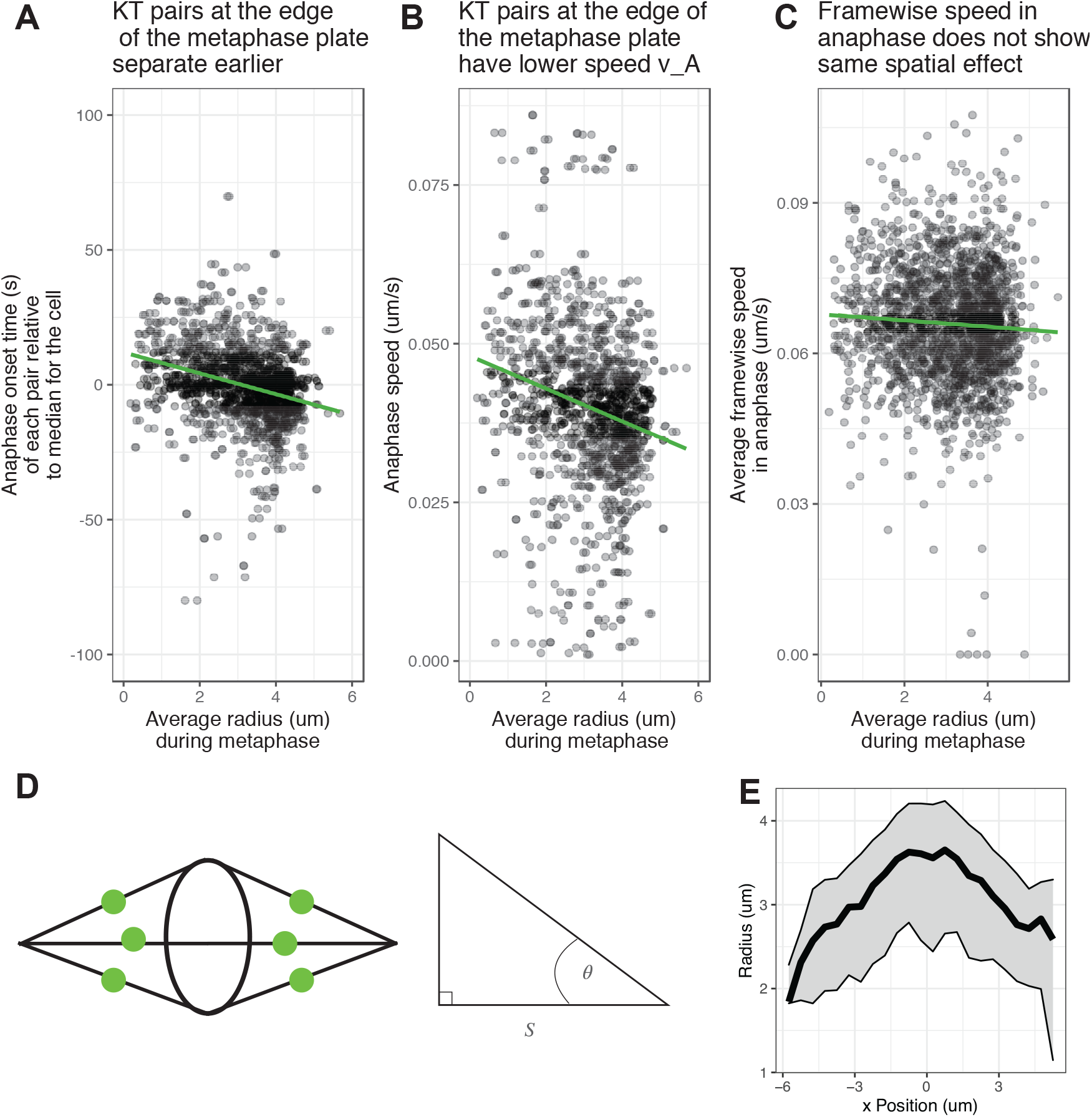
Anaphase onset occurs earlier at the edge of the metaphase plate. (A) Scatter plot of anaphase onset times relative to the median anaphase onset time in that cell versus the average radius of the kinetochore pair within the the metaphase plate. (B) Scatter plot of anaphase speed, *υ*_*A*_, versus the average radial position of of the kinetochore pair within the metaphase plate. (C) Scatter plot of the speed of kinetochore pairs in anaphase calculated framewise in 3D versus the average radial position of the kinetochore pair within the metaphase plate. For (A), (B) and (C), green lines indicate a linear fit to the data. (D) Schematic diagram indicating the metaphase plate and mitotic spindle (black lines) approximated via two cones, along with segregating kinetochores (green). The triangle depicted corresponds to the triangle between a kinetochore with average position at the centre of the metaphase plate, a kinetochore with average position at the edge of the plate, and one of the spindle poles. (E) Average radial location of kinetochores during anaphase for a given position on the *x* axis perpendicular to the metaphase plate. Solid black line shows median, with grey region showing 25% and 75% percentiles.

If we assume that both the kinetochore at the centre, and the kinetochore at the edge begin to segregate at the same time and move at the same speed, then the kinetochore at the edge takes approximately

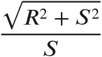

times longer to reach the pole. For realistic values of *R* = 4 *µ*m and *S* = 9 *µ*m (see Fig. 11E), then we find that the transit time for the kinetochore at the edge is 9% longer. Over the timescale of anaphase A of around 5 mins (***Su et al., 2016***), this gives a similar timescale to the difference in segregation times observed between kinetochore pairs at the edge and the centre of the metaphase plate. This suggests that anaphase onset triggers earlier at the periphery to coordinate chromosome separation.

### Mechanical forces at anaphase onset and force maturation in metaphase

From our annotated sister pair trajectories we are able to unlock a number of dynamic and mechanical questions. Using inferred parameters and K-fibre polymerisation states sampled from the posterior distribution, we computed the expected forces acting on a kinetochore throughout time (recall forces are quantified as speeds by dividing through by the unknown drag force). Averaging over kinetochore pairs reveals a force profile around anaphase onset as shown in Fig. 12A. The largest force in metaphase is the force due to polymerisation/depolymerisation of the K-fibre, with a smaller contribution from the centromeric chromatin spring and polar ejection force, a similar pattern to that seen in Hela cells (***Armond et al., 2015***). The net force (Fig. 12B) increases immediately after anaphase onset; during metaphase forces are in opposing directions.

**Figure 12.**
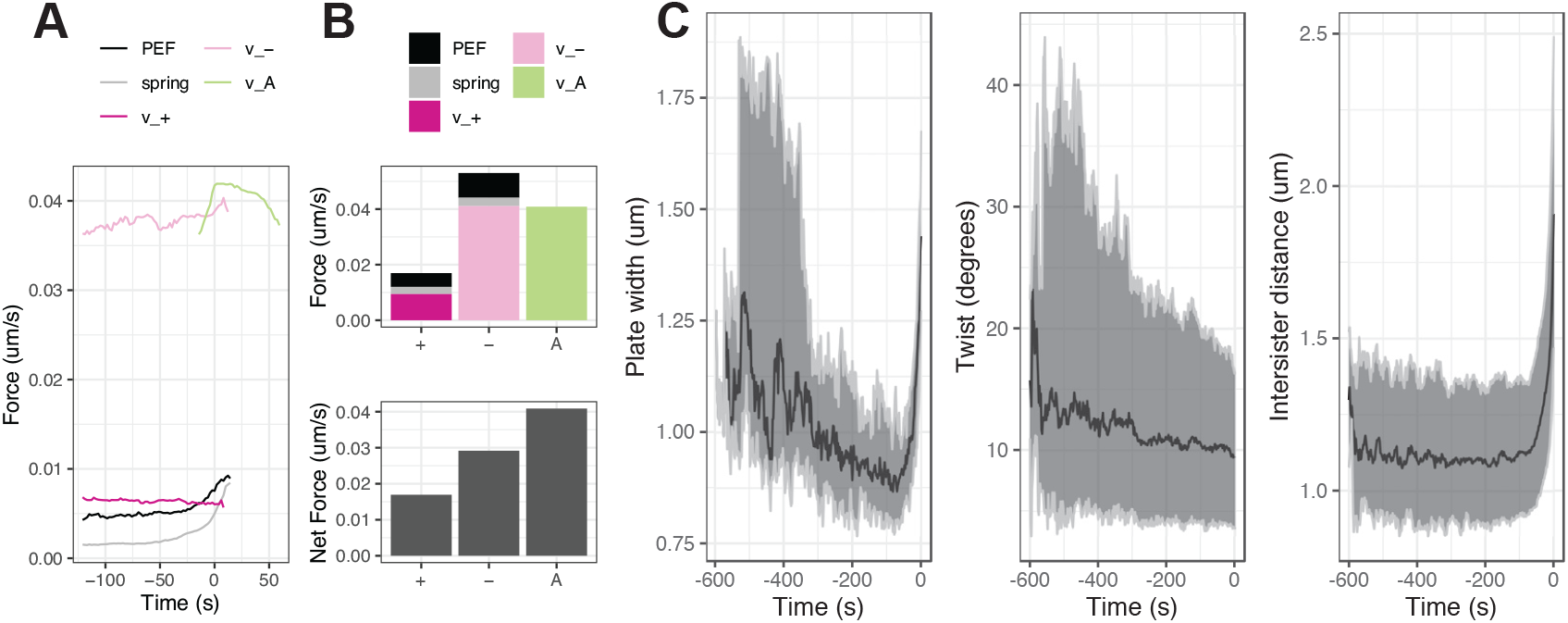
Force profiles at anaphase onset and kinetochore maturation characterisation during metaphase (A) Average force profile over time around anaphase onset across the population of kinetochores. Units of force terms are given as [um/s] as explained in Appendix 2. (B) Relative contributions of forces averaged over time shown as a bar plot (top) for K-fibres in polymerizing (+), depolymerizing (-) or anaphase (A) states, and total net force contributions in each of these states. (C) Changes over time during metaphase of the width of the metaphase plate (left), twist angle between the normal to the metaphase plate and the sister-sister vector (middle), and 3D intersister distance (right). Black line shows the median and grey region shows the 2.5% and 97.5% percentiles; data from *n* = 684 kinetochore pairs from *N* = 25 cells. The units of the inferred forces are [um/*s*] rather than [g um/*s*^2^] due to rescaling by the unknown drag coefficient.

Over the course of metaphase chromosome dynamics change, specifically there is an increase in centromere stiffness (***Harasymiw et al., 2019***) and a reduction of oscillation amplitudes (***Jaqaman et al., 2010***). By aligning trajectories of kinetochores to the median anaphase onset time of a cell, we can quantify these changes over time in higher resolution relative to a fixed reference point (anaphase onset). The metaphase plate becomes thinner over time (Fig. 12C, left) which is predominantly due to a decrease in the oscillation amplitude. The twist angle between the sister-sister axis and the normal to the metaphase plate reduces over time (Fig. 12C, middle) such that sisters become more aligned to the metaphase plate normal (which is likely to be aligned to the spindle axis (***McIntosh and Landis, 1971***)), suggesting a mechanical stiffening of the K-fibre-kinetochore attachment making it more rigid to torque (increased resistance to twist). The intersister distance between a kinetochore pair reduces slightly in metaphase until it begins to increase sharply as some kinetochore pairs separate (Fig. 12C, right).

## Discussion

In this work, we provide an in depth quantitative analysis of kinetochore dynamics in metaphase and anaphase A in untransformed human cells (RPE1). This required development of computational tools to fully utilise the high spatio-temporal resolution and quality achievable with LLSM. This included improvements in tracking software (KiT), development of a new metaphase-anaphase model and associated Bayesian inference algorithms, both for inference on data for single sister pairs and all the (tracked) sister pairs in a cell. We achieved near-complete kinetochore tracking, performed a detailed state and event annotation of kinetochore trajectories, analysed the heterogeneity of sister dynamics between and within cells and demonstrated that kinetochore dynamics are coordinated in time and space over the metaphase plate. We reveal that there is high variability of kinetochore behaviour within a cell, with position in the metaphase plate being a major determinant of behaviour, and that interactions between kinetochore pairs are important in governing metaphase dynamics and, potentially, the transition to anaphase. This implies that it is essential to consider chromosome dynamics in the context of the cell, accounting for the behaviour of neighbouring chromosomes and variations in mechanical properties throughout the spindle, rather than analysis of single kinetochore pairs in isolation.

We have demonstrated near-complete kinetochore tracking at high temporal resolution over long timescales from prometaphase to late anaphase. Two key factors enabled this. Firstly, we utilised LLSM (***Chen et al., 2014***) that uses an ultra thin sheet of light to limit effects of phototoxicity and photobleaching, thus allowing imaging over long timescales at high temporal resolution (***Sen et al., 2021***). Secondly, we made algorithmic improvements to our previous tracking algorithm (***Armond et al., 2016***), foremost among these being a change to the objective function in the adaptive detection step to include biological information about the expected number of kinetochores in an image.

We developed a mechanical switching model of chromosome dynamics in metaphase and anaphase, including dynamics at the metaphase-anaphase transition. To capture rare reversal events in anaphase, we extended this model from individual kinetochore pairs to a model of the kinetochore population of a cell, fitted to experimental data using a hierarchical framework whereby switching rate parameters are shared between kinetochore pairs in the same cell. Our model captures key forces and events, although it ignores the complex dynamics of K-fibres and the spindle. This simplicity ensures that all parameters of this model can be inferred from experimental trajectory data from a single cell (except for the natural spring length, see Methods). This would not be possible with a highly detailed molecular model with large numbers of parameters, since trajectory data is not informative on detailed molecular aspects governing the dynamics. Our model thus ignores some biological phenomena. Further, model parameters are assumed constant in time over the course of metaphase; however, mechanical maturation is observed during this time period, see Fig. 12 and ***Harasymiw et al. (2019***). Future extensions of the model could address such factors, along with directly modelling interactions between kinetochore pairs. In the hierarchical reversals model, both kinetochore sisters are assumed to be in the same anaphase/reversal state; a model with independent reversals for each kinetochore sister could also be considered.

We demonstrated significant heterogeneity in kinetochore dynamics within a cell; in fact the variation between kinetochore pairs in a cell is greater than variation between different cells (under the same experimental conditions). Previous work on HeLa and PtK1 cells demonstrated that biophysical parameters for the PEF (***Civelekoglu-Scholey et al., 2013***), *α*, the K-fibre forces, *υ*_−_, *υ*_+_, and diffusive noise *τ* vary spatially across the metaphase plate (***Armond et al., 2015***), with larger amplitude oscillations and lower noise kinetochores typically closer to the centre of the metaphase plate and noisier, lower amplitude oscillations at the edge of the metaphase plate (***Armond et al., 2015***; ***Civelekoglu-Scholey et al., 2013***). This is also true for RPE1 cells (Table 1), whilst we have further demonstrated that spatial variation is present in the switching events, Fig. 7. This suggested that location in the spindle strongly affects kinetochore behaviour, which may explain the higher variation within cells than between cells for certain parameters. This turned out to be the case; once radial position is accounted for variation between cells is comparable to variation within cells for the PEF parameter, *α*, (Fig. 6). In addition to metaphase plate position, there may be additional factors contributing to this variation between kinetochore pairs in a cell. For instance, dynamics may depend on specific chromosome properties, including size. This is suggested by the mis-segregation bias for certain chromosomes (***Worrall et al., 2018***), and for large kinetochores (***Drpic et al., 2018***). Our analysis thus suggests that metaphase dynamics are robust to changes in the spindle environment, for instance robust to variation in K-fibre length and composition, our data in fact indicating substantial spindle variation with transverse distance from the spindle axis. Despite these variations, qualitatively similar dynamics occurs throughout the plate.

This robustness of metaphase oscillations is also evident in comparison between cell types. We found that the centromeric chromatin spring is more compliant in RPE1 cells than in HeLa cells (***Armond et al., 2015***), corresponding to lower spring constant, *κ*, which is consistent with reduced breathing between sister kinetochores in HeLa cells compared to that in RPE1 cells. Whether stiffer centromeric chromatic springs is typical for cancer cells is unknown, but could explain disrupted centromere mechanical maturation (***Harasymiw et al., 2019***) in aneuploid cell lines, and recent observations of attenuated chromosome oscillations in cancer cell lines ***Iemura et al. (2021***). Similarly, the K-fibres forces *υ*_−_, *υ*_+_ have a larger difference in magnitude in RPE1 cells than in HeLa cells, with possible relevance to the increased kinetochore-microtubule dynamics reported in RPE1 cells relative to cancer cell lines (***Bakhoum et al., 2009***). This difference in K-fibre forces may be related to misattachment errors such as merotely; of note is that lazy kinetochores (***Sen et al., 2021***) in RPE1 cells have a higher value of *υ*_+_ than timely kinetochores, which adds further to the metaphase signature (including reduced intersister distance and impaired oscillation regulatiry) of lazy kinetochores reported in ***Sen et al. (2021***). In common with HeLa cells, we observed directional switching is typically, but not exclusively, initiated by the leading sister kinetochore.

We showed that kinetochores at the edge of the metaphase plate enter anaphase earlier, and have more directional switching events induced by the trailing kinetochore (and fewer induced by the leading kinetochore). One interpretation of the earlier onset of anaphase at the edge of the metaphase plate could be that in order to arrive near the spindle poles at the end of anaphase A in a synchronous manner, kinetochore pairs at the edge of the metaphase plate must segregate earlier because they have further to travel. This is consistent with calculations based on the simplified mitotic spindle geometry shown in Fig. 11E, and may potentially be due to separase (part of a positive feedback loop that increases the abruptness of anaphase (***Holt et al., 2008***)) penetrating to the cohesins of peripheral chromosomes earlier.

An analysis of the coordination between kinetochore behaviour within a cell demonstrated spatiotemporal interactions, including local coordination of directional switching events, kinetochore state and anaphase onset. Such local coordination is consistent with previous results reporting distance-dependent coupling of movements of neighbouring kinetochores in HeLa and RPE1 cells (***Vladimirou et al., 2013***), and load distribution of forces anchoring a kinetochore within ∼2um laterally (***Elting et al., 2017***) in PtK2 cells. Interestingly, HeLa and RPE1 cells have similar size spindles but very different numbers of kinetochores which is likely to affect the strength of the local coupling; ***Vladimirou et al. (2013***) found stronger correlated movements in HeLa than in RPE1 cells.

It is known that even weak physical coupling between oscillators leads to synchronisation, an observation that goes back to Huygens in 1665 (***Willms et al., 2017***). Thus, interactions, or coupling between non-sister chromosomes would lead to oscillation synchronisation across the plate, akin to a beating drum, although microtubule dynamics are intrinsically noisy which would cause deterioration in the synchronisation. The spindle is surrounded by the endoplasmic reticulum (***Ferrandiz et al., 2021***) which likely increases the frictional and drag forces on the peripheral chromosomes, thus reducing the amplitude of those oscillations towards the periphery. Inter-chromosome interactions could arise through many mechanisms. The spindle is a visco-elastic anisotropic material (***Shimamoto et al., 2011***) with chromosomes moving within the spindle primarily along the spindle axis. Coupling between neighbouring chromosomes through this material would thus be expected. In fact, viscosity alone is known to induce hydrodynamic forces between moving objects through fluid movements. Thus, interactions that generate the space-time coordination observed in our data could be a result of spindle material properties (visco-elastic or hydrodynamic), or arise from direct interactions between kinetochore pairs (such as cross linking of K-fibres as proposed in ***Vladimirou et al. (2013***)), or between the chromosome arms, Ki67 acting as a surfacant to reduce adherence in mitosis (***Cuylen et al., 2016***). The simplest hypothesis is that inherent variation in local spindle architecture modulates mechanics locally at each point in metaphase plate, changes in spindle geometry resulting in variation of the PEF with metaphase plate position (***Armond et al., 2015***; ***Civelekoglu-Scholey et al., 2013***) and inducing kinetochore swivel (***Smith et al., 2016***), whilst material interactions (eg. drag) within the spindle environment and between chromosomes (***Cuylen et al., 2016***) couple neighbouring kinetochores, which generates coordination between neighbouring kinetochore movements. Finally, anaphase onset may be locally coordinated because of the above mechanisms, or potentially through spatial variation in separase activity across the plate. Thus we highlight how local interactions between kinetochores and feedback between spindle architecture and mechanics (***Elting et al., 2018***) can give rise to emergent properties of mitotic cells and ensure robustness of chromosome dynamics.

This study provides unprecedented detail and analysis of the behaviour of the complement of chromosomes across the metaphase-anaphase transition in non-transformed human RPE1 cells. This comprehensive characterisation should prove invaluable when comparing cell phenotypes (***Pargett and Umulis, 2013***) and assessing changes to chromosome dynamics upon perturbations, be it genetic or through application of drugs. Thus, our methodology will allow mechanistic hypotheses to be evaluated with high precision. This work sets the stage for future work to quantitatively analyse the whole of mitosis at the cell level, from nuclear envelope breakdown through to segregation of chromosomes.

## Methods and Materials

### Code

Code used to produce the results reported in this work is available at www.github.com/shug3502/MitosisModels/.

### Cell culture and generation of cell lines

Immortalized (hTERT) diploid human retinal pigment epithelial (RPE1) cell line (MC191), expressing endogenously tagged Ndc80-eGFP, was generated by CRISPR-Cas9 gene editing ***Roscioli et al. (2020***). hTERT-RPE1 cells were grown in DMEM/F-12 medium containing 10% fetal bovine serum (FBS), 2 mM L-glutamine, 100 U/ml penicillin and 100 mg/ml streptomycin (full growth medium); and were maintained at 37°C with 5% CO_2_ in a humidified incubator.

### Live cell imaging by lattice light sheet microscope

The lattice light sheet microscope (LLSM) ***Chen et al. (2014***) used in this study was manufactured by 3i (https://www.intelligent-imaging.com). Cells were seeded on 5 mm radius glass coverslips one day before imaging. On the imaging day, each coverslip was transferred to the LLSM bath filled with CO_2_-independent L15 medium, where live imaging takes place. All imaged cells entered anaphase, which is a suitable proxy for a lack of phototoxicity effects ***Jaqaman et al. (2010***). The LLSM light path was aligned at the beginning of every imaging session by performing beam alignment, dye alignment and bead alignment, followed by the acquisition of a bead image (at 488 nm channel) for measuring the experimental point spread function (PSF). This PSF image is later used for the deconvolution of images. 3D time-lapse images (movies) of Ndc80-eGFP were acquired at 488nm channel using 1% laser power, 20 ms exposure time/z-plane, 75 z-planes, 307 nm z-step and 0.5 s laser off time, which results in 2 s/z-stack time/frame. Acquired movies were de-skewed and cropped in XYZ and time, using Slidebook software in order to reduce the file size. Cropped movies were then saved as OME-TIFF files in ImageJ.

### Tracking

Kinetochore tracking is performed using the software package KiT (***Armond et al., 2016***) v2.4.0. The tracking algorithm proceeds by detecting candidate spots via an adaptive threshold method to set a threshold on the image histogram. Candidate spot locations are refined by fitting a Gaussian mixture model. Spot locations are linked between frames by solving a linear assignment problem, with motion propagation via a Kalman filter. Tracked kinetochores are paired by solving another linear assignment problem.

Code to perform kinetochore tracking is available from https://github.com/cmcb-warwick/KiT, and this software includes a graphical user interface (GUI) for ease of use.

### Likelihood calculation

To compute the likelihood for this model we apply the forward algorithm and marginalize out the discrete hidden states. Let 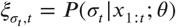 be the probability of being in state *σ*_*t*_ at time *t* given all the data up to that time. Suppose that we define the likelihood contribution of an observation, *x*_*t*_, as 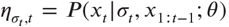. Notably, for the SDE model considered here, the likelihood contribution of an observation, *x*_*t*_, depends only on the previous observation and not the entire history:

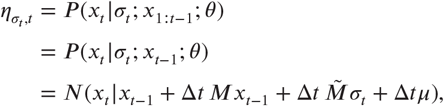

where *N*(·| ·) is the normal density function, and 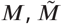, and *µ* are derived from the linear SDE in Eq. (1), and given by

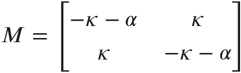

and

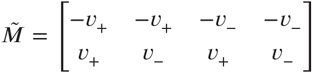

and

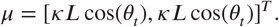

In this notation, the state variable 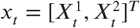 represents the positions of both sisters in a pair at time *t* and the state *σ*_*t*_ ∈ {[1, 0, 0, 0]^*T*^, [0, 1, 0, 0]^*T*^, [0, 0, 1, 0]^*T*^, [0, 0, 0, 1]^*T*^} corresponding to the states {++, +−, −+, −−} for the sister kinetochore pair at time *t*.

Replacing these in the expressions above gives

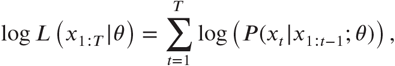

where

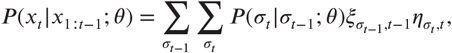

and hidden state probabilities are determined iteratively via:

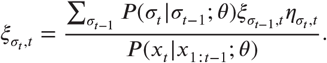

### Sampling from the distribution of the hidden states

The expression required from the forward-backward algorithm is achieved by combining the results of the forward and backward algorithms as follows:

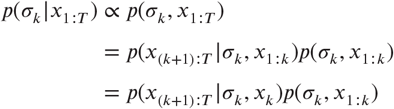

due to the graphical structure of the model (as in Fig. **??**). Including the normalizing term, we have

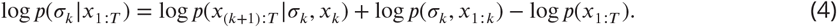

### Forward algorithm

The following derivation proceeds by summing over all possible values of the hidden state variable *σ*_*k*−1_, using the definition of conditional probability, and the graphical structure of the model.

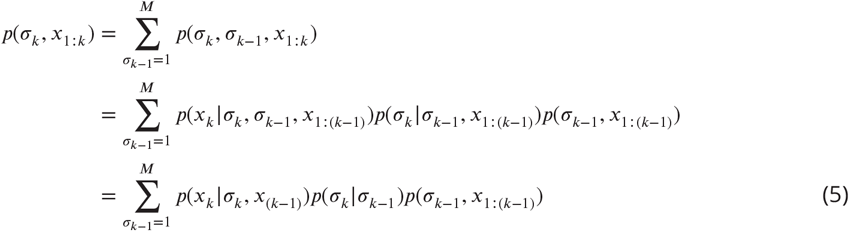

where *p*(*x*_*k*_ | *σ*_*k*_, *x*_(*k*−1)_) is available via the SDE, *p*(*σ*_*k*_| *σ*_*k*−1_) is given via the transition matrix, and *p*(*σ*_*k*−1_, *x*_1:(*k*−1)_) is equivalent to the term on the left hand side (LHS) for a different value of *k*. Applying this result iteratively forward in time from an initial condition allows us to solve for *p*(*σ*_*k*_, *x*_1*:k*_).

### Backward algorithm

Similar steps are applied to derive a recursion for the backward algorithm.

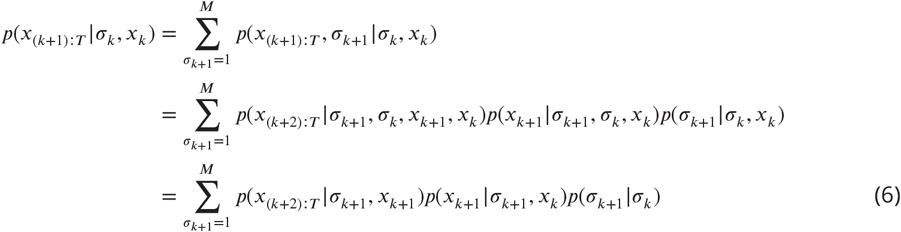

where *p*(*x*_(*k*+2):*T*_ | *σ*_*k*+1_, *x*_*k*+1_) is an iterated version of the LHS, *p*(*x*_*k*+1_ | *σ*_*k*+1_, *x*_*k*_) is available via the SDE, and *p*(*σ*_*k*+1_ | *σ*_*k*_) is given via the transition matrix. We apply this iteratively from *k* = *T* − 1, …, 1.

### Backward sampling

In order to make statements about switching events, we need to consider sequences of states forming a pattern corresponding to coherent switches from one coherent state to another via an intermediate state. To address this, we sample from the full hidden state sequence given all the data, and assess this for switches.

To enable the sampling, we observe that the pairwise marginal distribution can be expressed as

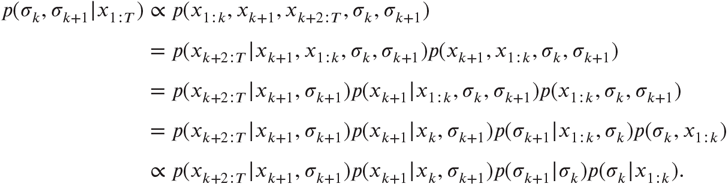

This expression contains known terms and terms that can be computed via the forward and backward algorithms. Furthermore, when *σ*_*k*+1_ is known and we are considering how to sample back in time through the hidden states, we have that

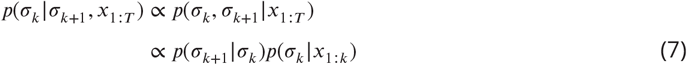

since the other terms in the expression above do not depend on *σ*_*k*_.

The strategy for the backward sampling algorithm is therefore to sample initially from *p*(*σ*_*T*_| *x*_1*:T*_), which is available from the forward algorithm, and subsequently to simulate backward in time from *T* to 1 via the conditional distribution given in Eq (7).

### Prior distributions

We impose broad prior distributions on the parameters of the biophysical model, as shown in Supp. Table 1. For the natural length of the spring, *L*, we impose an informative prior based on an additional nocodazole washout experiment (nocodazole interferes with polymerization of microtubules). This avoids an unidentifiabilty in the model as in ***Armond et al. (2015***). Additionally, we use an informative prior for the time of anaphase, *t*_*A*_, based on first fitting a changepoint model (with a uniform prior on *t*_*A*_) to get an initial estimate for anaphase onset to guide the biophysical model and avoid exploring parameter space corresponding to pathological behaviour such as anaphase at the start or end of movies.

### Convergence diagnostics

Convergence and mixing of MCMC chains is assessed via the Gelman-Rubin 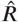 statistic ***Gelman and Rubin*** (***1992***); ***Vehtari et al. (2021***) using only results where 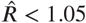 for all parameters.

Of *N* = 58 cells tracked, MCMC chains were run successfully for *N* = 32 cells. Where MCMC chains failed to run this was due either to poor tracking results in that cell (insufficient tracked kinetochore pairs) or long time series such that the MCMC chains progressed extremely slowly (failed to find the typical set). Of the *n* = 885 kinetochore pairs across 32 cells where MCMC chains ran successfully, MCMC chains from *n* = 201 kinetochore pairs failed to converge as assessed by the 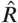 statistic, leaving estimates from *n* = 684 kinetochore pairs. Of the 32 cells with MCMC results, chains failed to converge on any kinetochore pairs for 7 cells as assessed by the 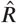 statistic. Thus results are reported for *n* = 684 kinetochore pairs from *N* = 25 cells.

### Changepoint model

A simple changepoint model (***Armond et al., 2019***) to assess the time of anaphase assumes that the intersister distance between a kinetochore pair is constant in metaphase, and increases linearly in time during anaphase. If *d*_*t*_ is the 1D intersister distance (in the *x* direction perpendicular to the metaphase plate) between kinetochore sisters, then at time point *t*_*i*_

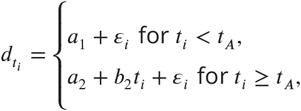

with the condition that *a*_1_ = *a*_2_ + *b*_2_*t*_*A*_. Since this model is simpler than the biophysical model, a uniform prior can be used for *t*_*A*_ and weakly informative priors for other parameters.

### Hawkes process model

The Hawkes process is a point process similar to an inhomogeneous Poisson process, with the key difference that the intensity depends on past events. The conditional intensity given the history of the process up to time *t* with switching events at times *t*_1_, …, *t*_*n*_ is

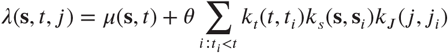

where we have kernels in time, *k*_*t*_(*t, t*_*i*_), in space, *k*_*s*_(***s, s***_*i*_), and between kinetochore pairs, *k*_*J*_ (*j, j*_*i*_). We take an exponential kernel in time

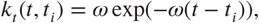

with timescale 1/*ω*, along with a Gaussian kernel in space

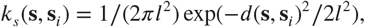

where *l* is the length scale and *d*(***s, s***_*i*_) is the Euclidean distance between ***s*** and ***s***_*i*_. The kernel between kinetochore pairs ensures that we consider only interactions between switches of different pairs

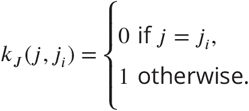

We assume that the background rate is constant in time and space such that *μ*(***s***, *t*) = *μ*_0_.

To compute the likelihood for the Hawkes process model, we follow methods used in ***Loeffler and Flaxman*** (***2018***). The likelihood of the Hawkes process model is as for an inhomogeneous Poisson process, and is given by

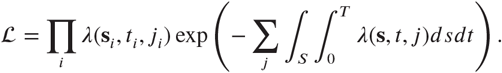

The log likelihood can be written as

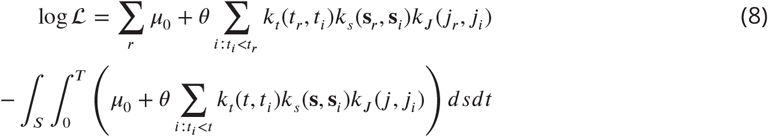

To simplify this expression, note that the final part of the integral term equates to the total number of events in the time interval considered

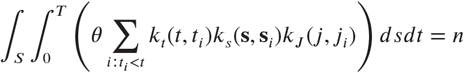

assuming the kernels used are normalized; this term is therfore constant with respect to the parameters of the Hawkes process which simplifies the likelihood computation.

We implement this model via the log likelihood computation above in Stan (***Carpenter et al., 2017***) and run 4 MCMC for 1000 iterations allowing us to sample from the posterior distribution via Hamiltonian Monte Carlo.

### Simulation from 4 state Markov model

To consider fluctuations in the proportion of kinetochore pairs in each state, we consider forward simulations from the Markov model for the hidden states in the anaphase model (as in Fig. 2C), but only including the metaphase states, (++,+-,-+,–). The transition matrix thus depends on *p*_coh_ and *p*_icoh_, the framewise probability of switching out of a given state for an individual kinetochore sister. Independent simulations are performed for *N* = 46 kinetochore pairs (corresponding to the number of kinetochore pairs in untransformed human cells) and averaged to give the proportion of pairs in a given state.

## Acknowledgments

JUH, OS, NJB and ADM are supported by BBSRC (BB/R009503/1). ADM is also supported by a Wellcome Senior Investigator Award (grant 106151/Z/14/Z). The Lattice Light Sheet Microscope Facility was established at Warwick with a Wellcome Trust Multi-user Equipment grant to ADM (grant 208384/Z/17/Z). We thank the Computational and Microscopy Development Unit (CAMDU) for support with lattice light sheet microscopy.

## Additional Information

### Author contributions

Jonathan U Harrison, Conceptualization, Methodology, Software, Formal Analysis, Writing - Original Draft, Writing - Review and Editing. Onur Sen, Conceptualization, Investigation, Design of Experiments and Data Collection, Writing - Review and Editing. Andrew D McAinsh, Conceptualization, Supervision, Project Administration, Funding Acquisition, Writing - Review and Editing. Nigel J Burroughs, Conceptualization, Methodology, Supervision, Project Administration, Funding Acquisition, Writing - Review and Editing.

## Appendix 1

### Supplementary Figures

**Appendix 1 Figure 1. Supplementary Figure 1.**
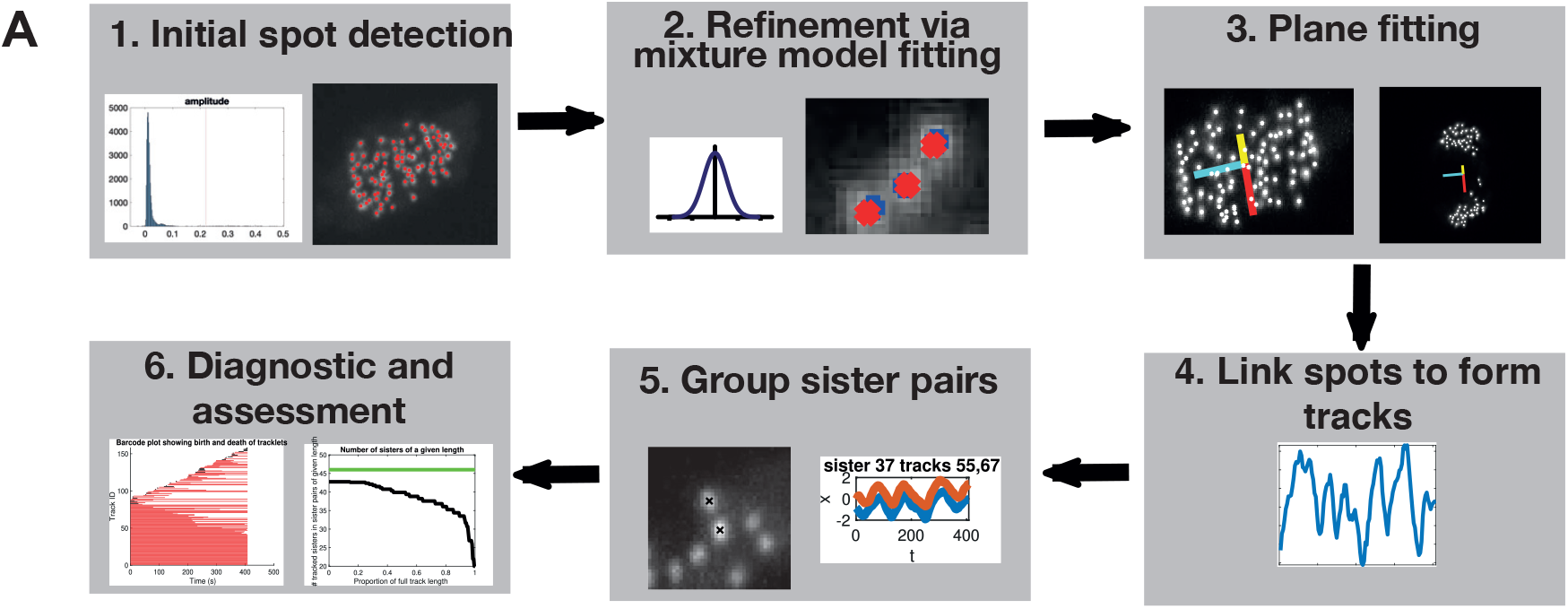
Tracking pipeline for near-complete tracking of kinetochores. Candidate spots are detected via an adaptive threshold technique (step 1). Spot locations are refined using a Gaussian mixture model (step 2). A plane is fitted to orientate a reference coordinate system with respect to the metaphase plate (step 3). Detected particles are linked between frames over time to form tracks (step 4). Kinetochore sister pairs are grouped based on metaphase dynamics (step 5). Tracks of kinetochore sister pairs can be assessed via diagnostic tools (step 6).

**Appendix 1 Figure 2. Supplementary Figure 2.**
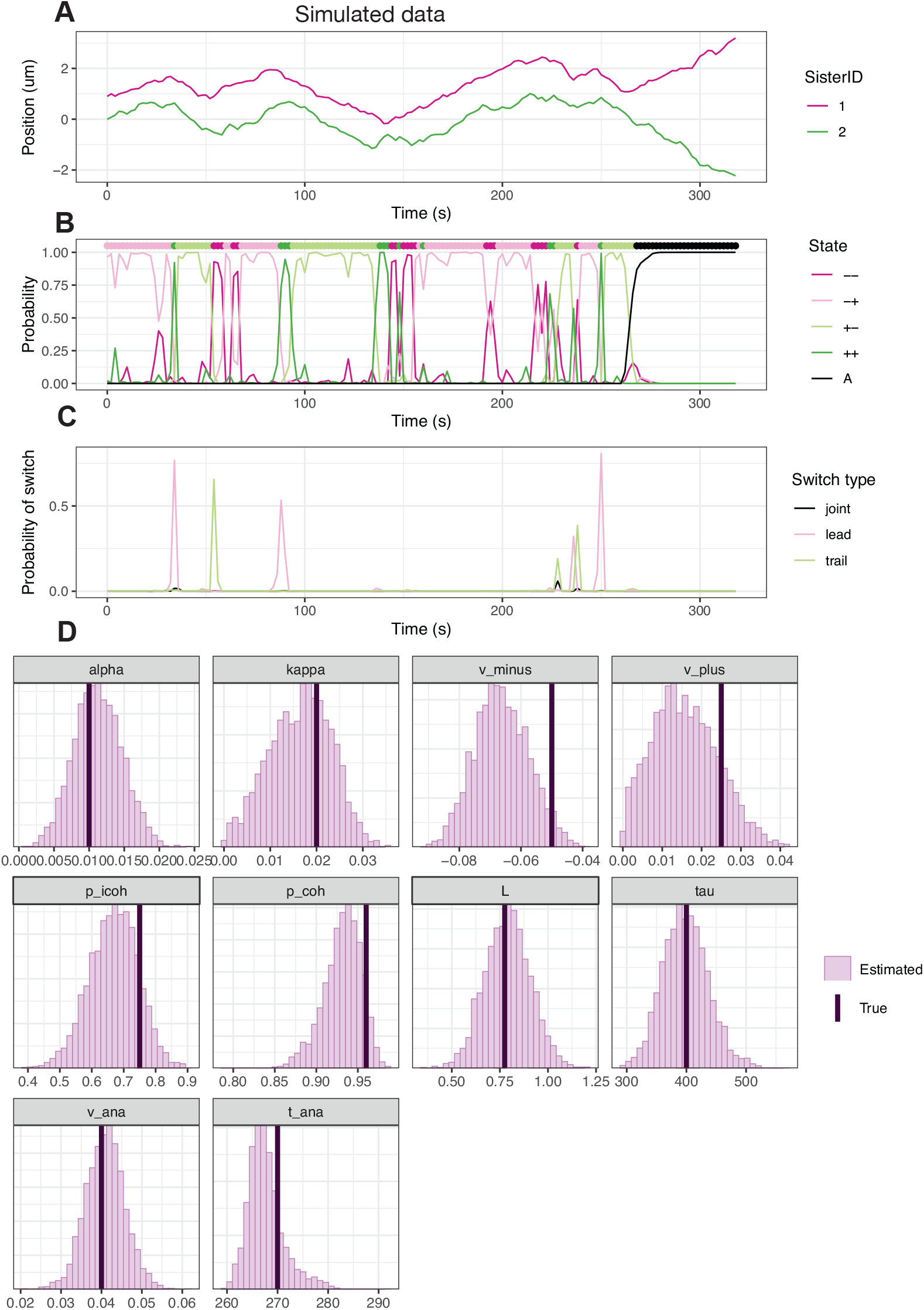
Estimation of parameters of the biophysical mode and annotation of hidden states on simulated data. (A) Simulated data from the anaphase model until a final time of 320 *s* (160 frames) with parameters: *τ* = 400, *α* = 0.01 *s*^−1^, *κ* = 0.02 *s*^−1^, *υ*_−_ = −0.05 *um/s, υ*_+_ = 0.025 *um/s, p*_icoh_ = 0.75, *p*_coh_ = 0.96, *L* = 0.775 *um, υ*_*A*_ = 0.04 *um/s, t*_*A*_ = 270 *s*, Δ*t* = 2 *s*. (B) Hidden microtubule attachment states in the simulated data and probabilities of each state as sampled from the posterior. Coloured points at the top of the plot indicate the true simulated hidden state, while the lines show the estimated probability of each state. (C) Switching probability based on patterns of hidden states sampled from the posterior distribution. Note that only switches from one coherent state (+− or −+) to the opposite coherent state, via an intermediate incoherent state (++ or −−) are considered. (D) Marginal posterior histograms for each of the model parameters. Vertical lines indicate the true parameters used to simulate the data. These lines lie within the posterior density indicating that the parameters used to simulate the data can be recovered.

**Appendix 1 Figure 3. Supplementary Figure 3.**
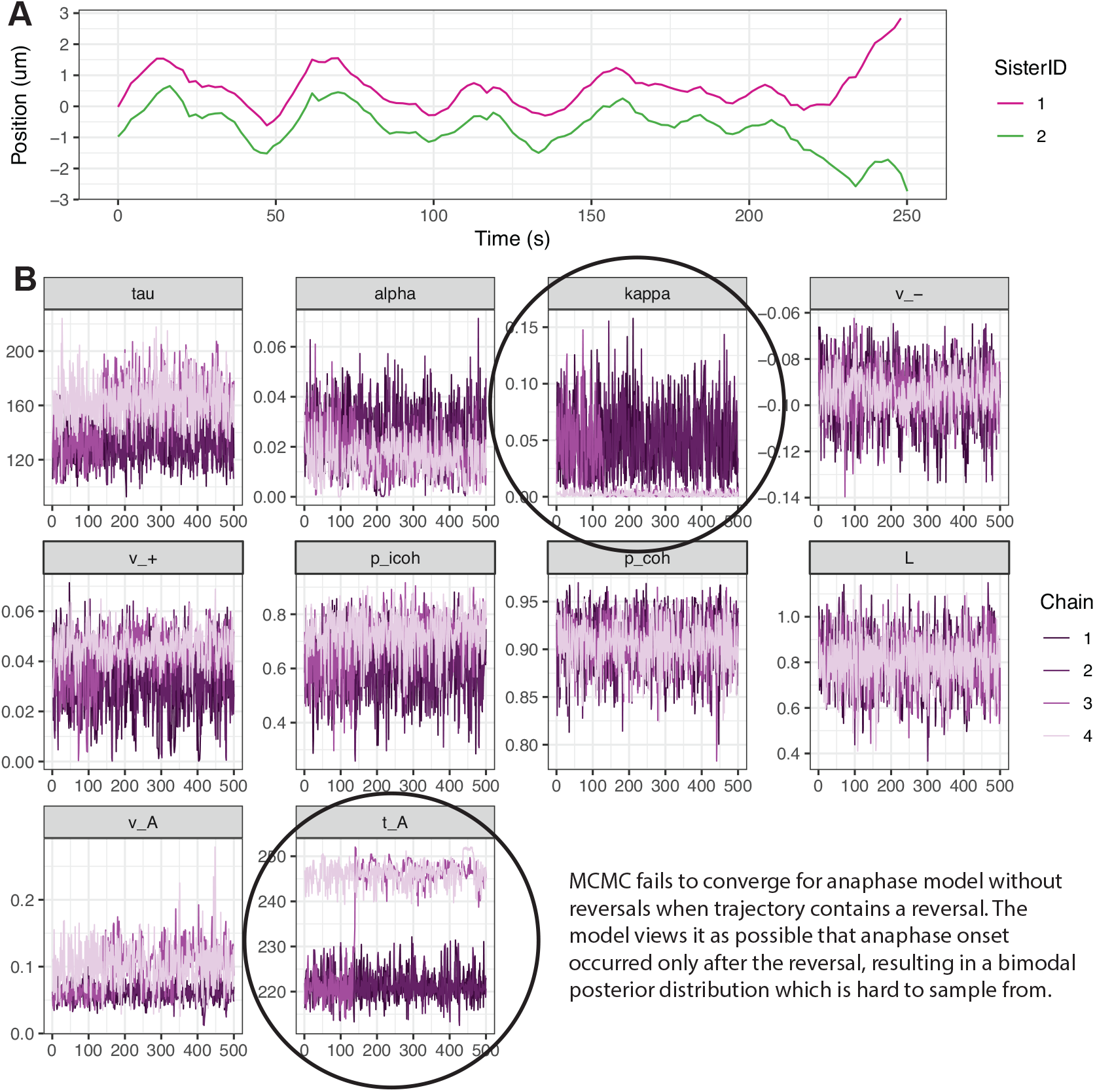
MCMC convergence issues for anaphase model when reversals are present in the data. (A) Kinetochore tracking data for a single kinetochore pair exhibiting a reversal away from the pole in anaphase (Sister 2 around 230s), before returning to poleward motion. (B) MCMC traceplots for 4 chains and all model parameters of the anaphase model. For parameters *κ* (spring constant) and *t*_*A*_ (time of anaphase onset) the chains have not mixed properly due to a bimodal posterior distribution. The posterior has one model for a normal or early anaphase onset when the kinetochore pair separates at around 220s, and another mode for a late anaphase onset at around 245s after the reversal.

**Appendix 1 Figure 4. Supplementary Figure 4.**
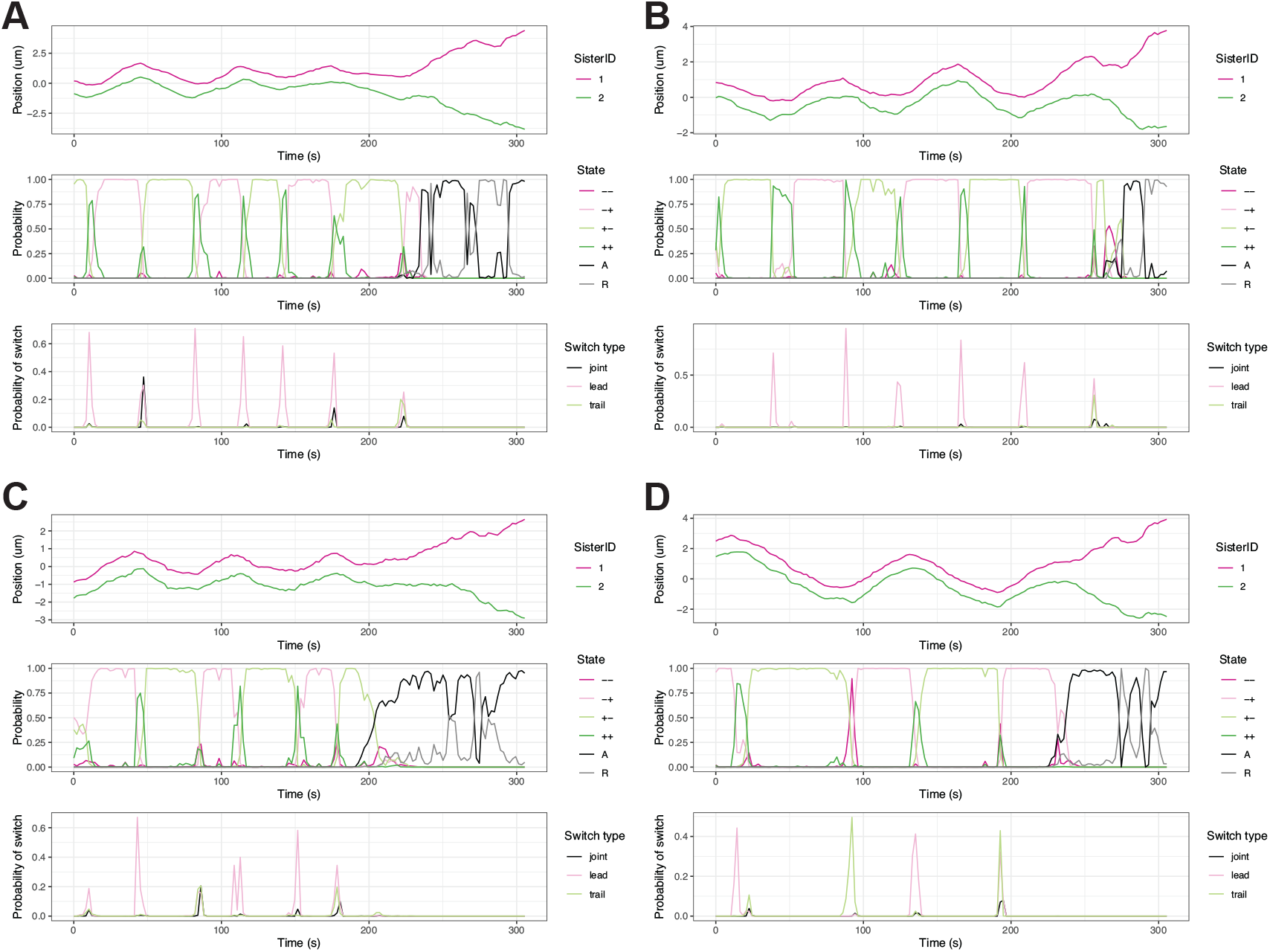
Example trajectories and inferred hidden states (and probability of a switch based on these states) using the hierarchical anaphase model with reversals for 4 different kinetochore pairs shown in (A), (B), (C), and (D). All four pairs are from the same cell shown in Figure 4.

**Appendix 1 Figure 5. Supplementary Figure 5.**
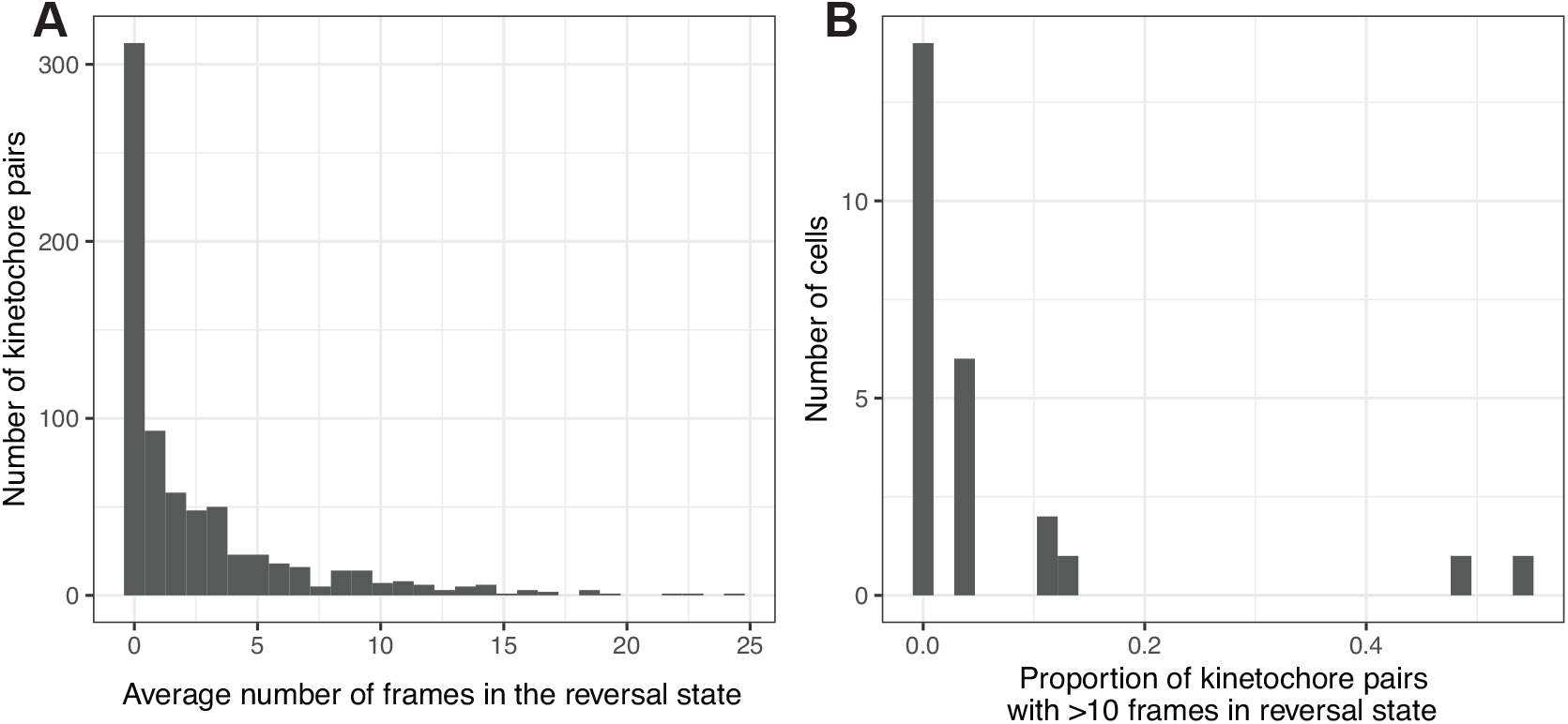
Reversals are rare events for most cells. (A) Histogram of the average number of frames spent in the reversal state over the population of kinetochore pairs. (B) Histogram of the proportion of kinetochore pairs in a cell that spent on average more than 10 frames in the reversal state, *R*. Results from *N* = 25 cells and *n* = 676 kinetochore pairs.

**Appendix 1 Figure 6. Supplementary Figure 6.**
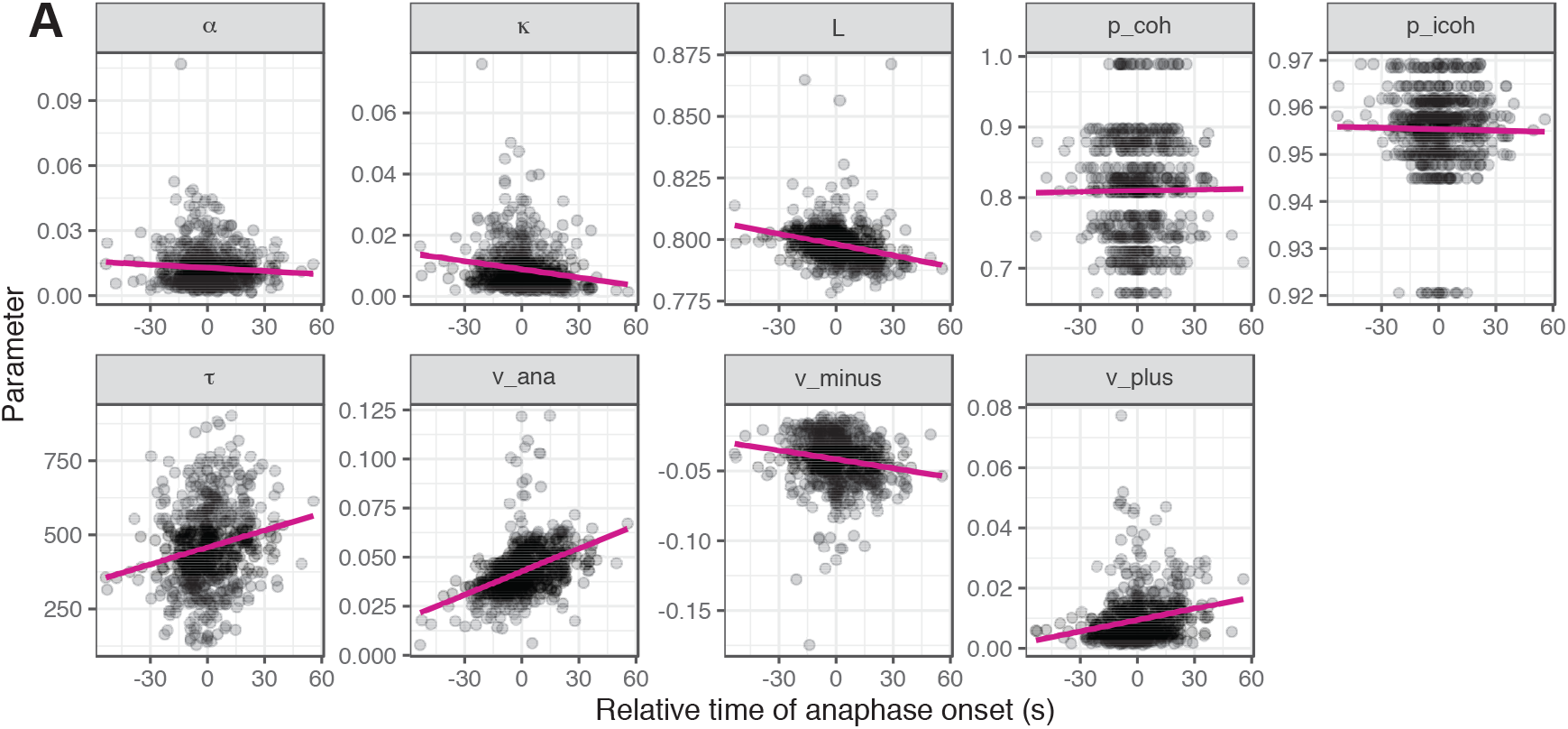
Correlations between relative time of anaphase onset and model parameters, based on median of estimated posterior distributions. Each point corresponds to a kinetochore pair, with *n* = 684 kinetochore pairs from *N* = 25 cells. Solid magenta line shows linear fit between time of anaphase onset (relative to the median over pairs in a cell) and the estimated median model parameters.

**Appendix 1 Figure 7. Supplementary Figure 7.**
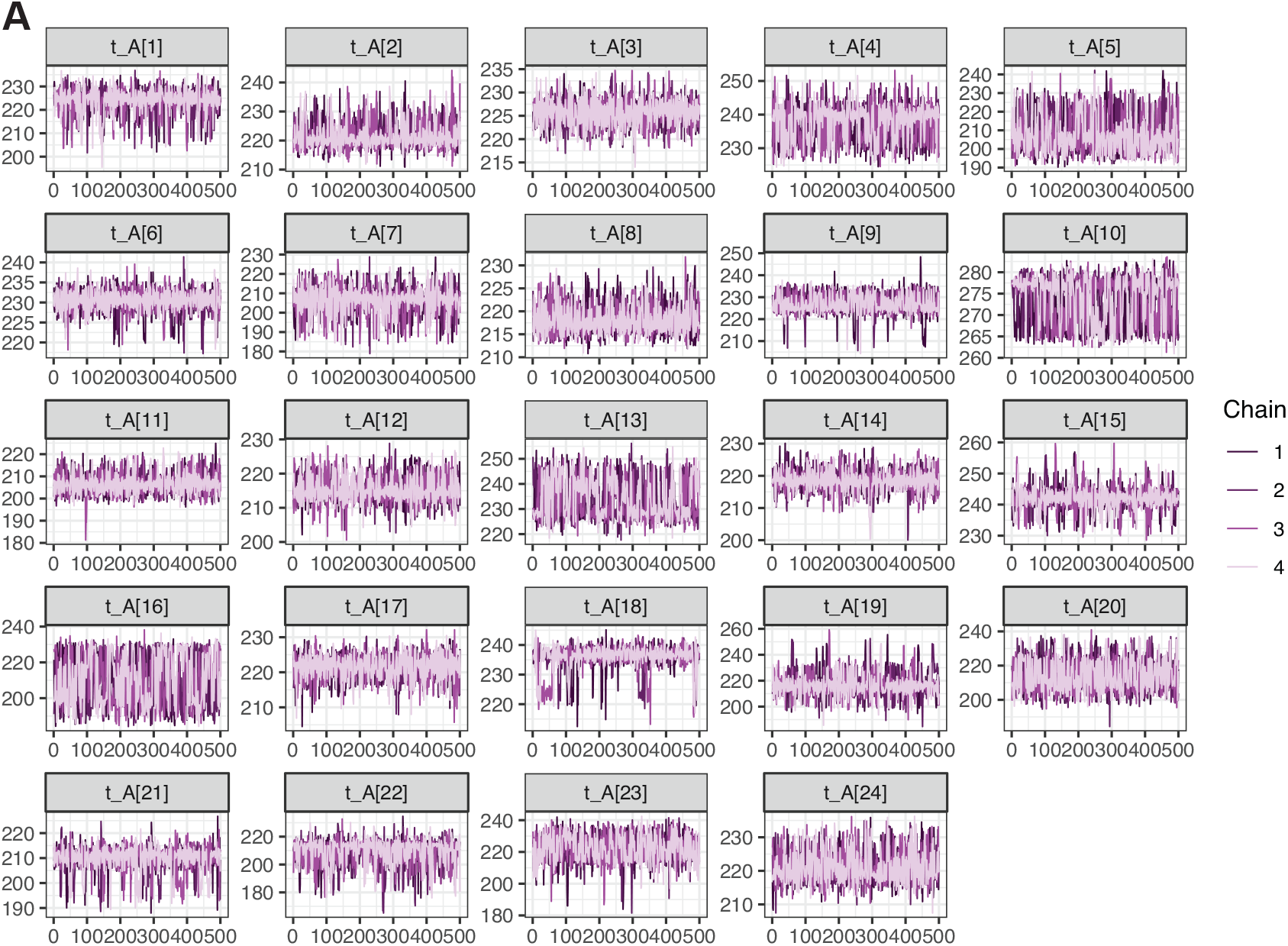
MCMC traceplots based on the anaphase model with reversals for the time of anaphase onset, *t*_*A*_, for each of the tracked kinetochore pairs in a cell. Four MCMC chains are run (see Methods) and are shown in different colours. The time of anaphase onset can have a bimodal distribution for some kinetochore pairs (eg. pairs 10 and 13 here) and thus checking traceplots for this parameter in particular can highlight problems with converged or mixing of MCMC chains. All pairs shown have 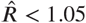 and including the reversal state, *R*, has helped avoid convergence and mixing issues present without this state.

**Appendix 1 Figure 8. Supplementary Figure 8.**
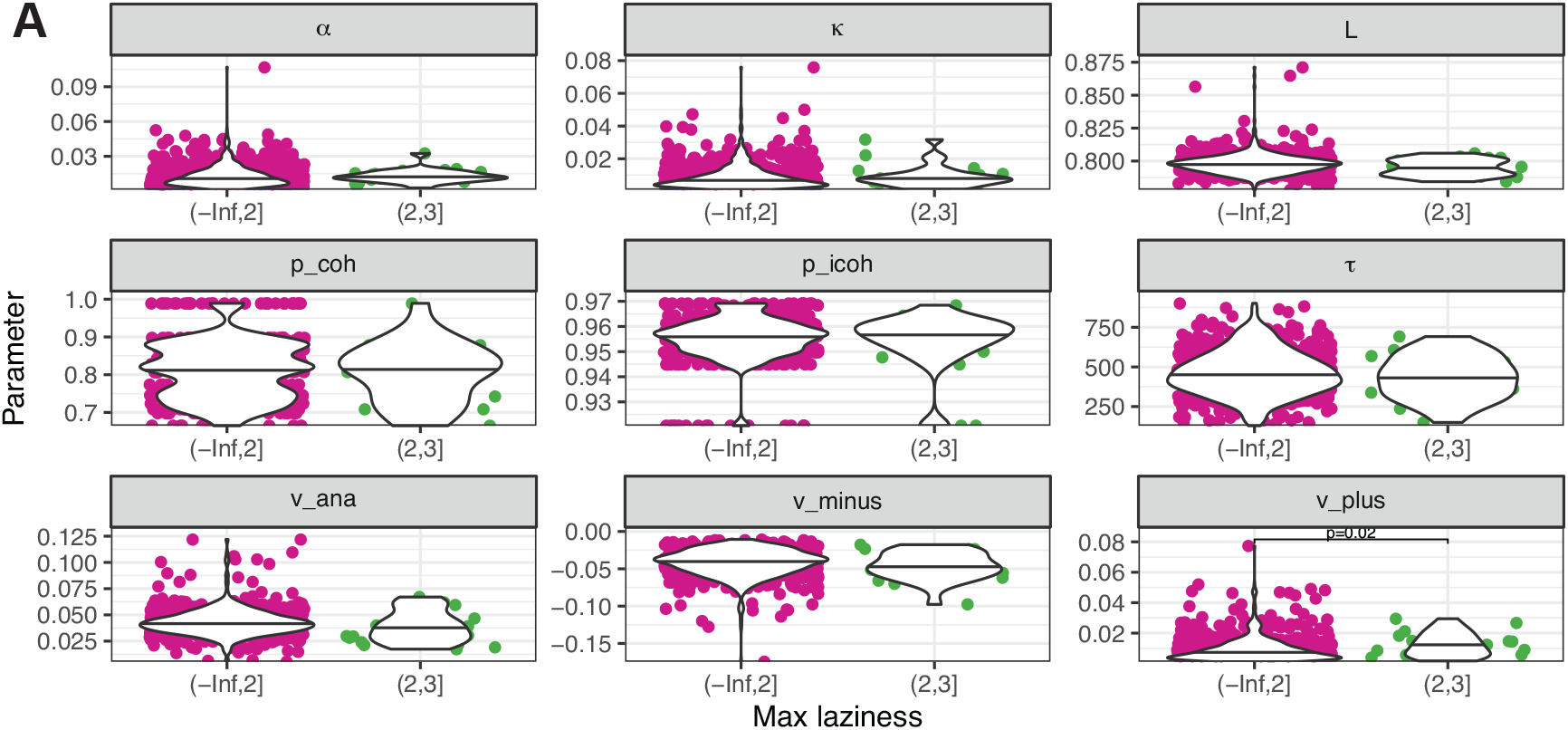
Distribution of inferred values for the biophysical parameters across kinetochore pairs with high laziness (>2) versus low laziness (<2). For definition of laziness, see ***Sen et al. (2021***). Results from *N* = 25 cells and *n* = 684 kinetochore pairs.

**Appendix 1 Figure 9. Supplementary Figure 9.**
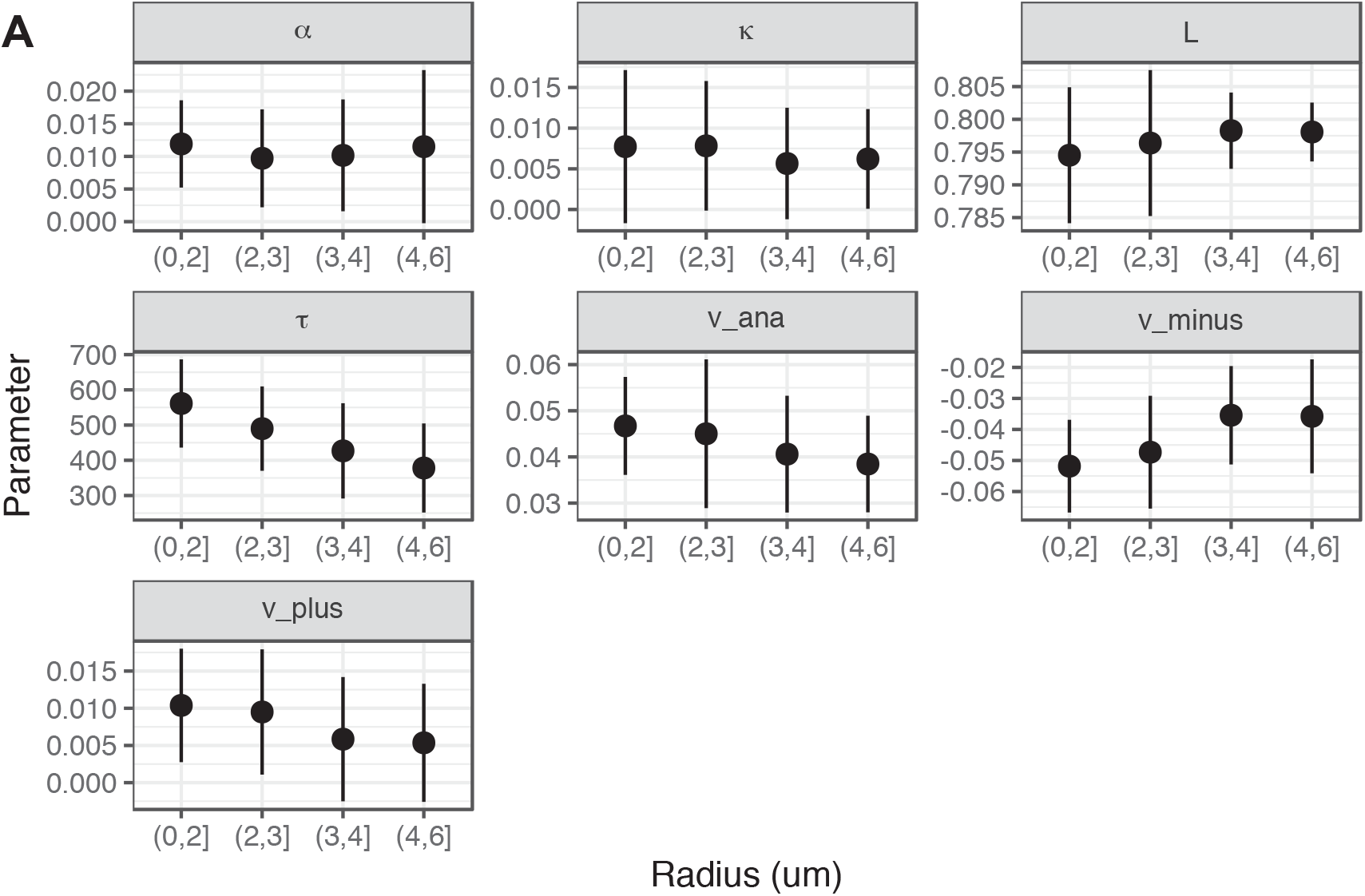
Dependence of biophysical parameters on radial position within the metaphase plate. Points indicate medians across *n* = 684 kinetochore pairs from *N* = 25 cells at a given radial location. Lines indicate median ± one standard deviation across the population (at a given radial location). Radial position is divided into (unequal sized) bins.

**Appendix 1 Figure 10. Supplementary Figure 10.**
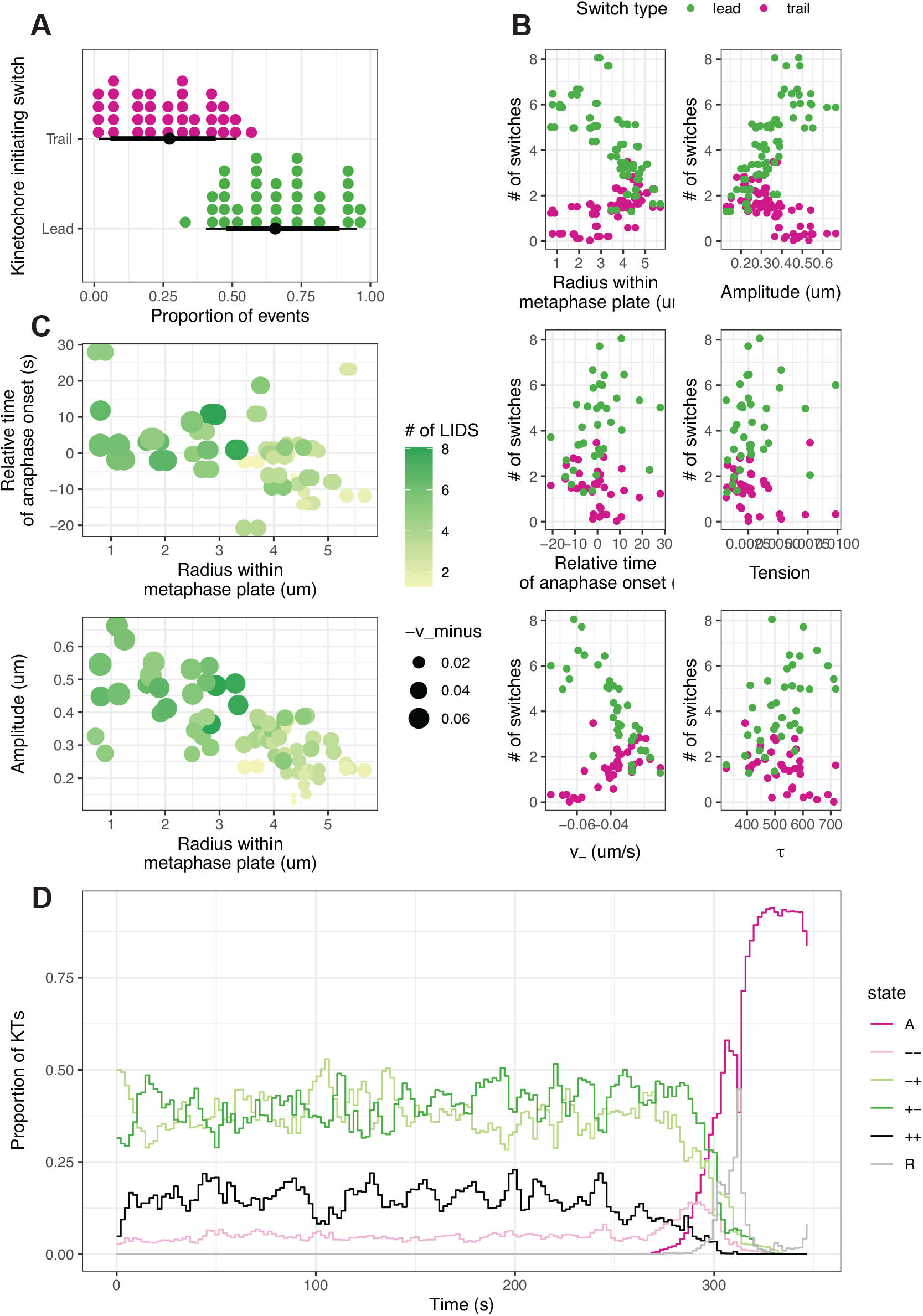
Directional switches of oscillating chromosomes vary across the metaphase plate. As for Figure 7, but for a different cell to indicate variability between cells. (A) Fraction of LIDS (green) and TIDS (pink) events as a proportion of the total number of switching events including joint switches. Each kinetochore pair gives rise to a LIDS and TIDS dot. (B) Relationship between the number of directional switches initiated by the leading (green) or trailing (pink) kinetochore sister, and other summary statistics describing the oscillatory dynamics. (C) Relationship between the number of LIDS events and other summary statistics indicating that many of these variables change together based on spatial position of kinetochore pairs within the metaphase plate. (D) Proportion of kinetochore sister pairs in a given hidden state at each time point.

**Appendix 1 Figure 11. Supplementary Figure 11.**
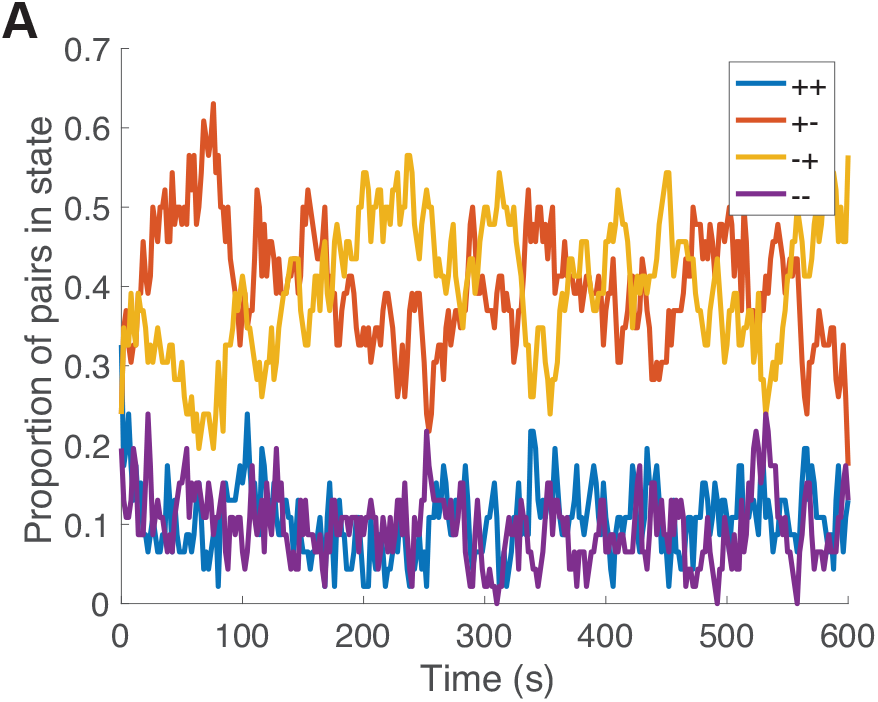
Simulating from a 4 state Markov model with spontaneous switching between states (see Methods) for *N* = 46 kinetochore pairs exhibits fluctuations in the proportion of kinetochore pairs in each state. Parameters used in simulation (*p*_coh_, *p*_icoh_) = (0.96, 0.83).

**Supplementary Table 1:** **Priors for the anaphase model and anaphase reversals model** where *t*\* is the median posterior marginal estimate for *t*_*A*_ using the changepoint model to provide an informative prior for *t*_*A*_.

| Parameter | Prior | Support |
| --- | --- | --- |
| $\alpha$ | $N(0.01, 0.1)$ | $[0, \infty)$ |
| $\kappa$ | $N(0.05, 0.1)$ | $[0, \infty)$ |
| $v_-$ | $N(-0.03, 0.1)$ | $(-\infty, 0]$ |
| $v_+$ | $N(0.03, 0.1)$ | $[0, \infty)$ |
| $p_{\text{icoh}}$ | $Beta(2, 1)$ | $(0, 1)$ |
| $p_{\text{coh}}$ | $Beta(2.5, 1)$ | $(0, 1)$ |
| $L$ | $N(0.79, 0.119)$ | $[0, \infty)$ |
| $\tau$ | $N(t^*, 14)$ | $[0, \infty)$ |

## Appendix 2

### Derivation of metaphase dynamics equations

To derive the metaphase dynamics in eq. (1), we proceed as follows. For kinetochore sister *j*, we apply force balance, ignoring inertial forces since the system is in a high viscous limit:

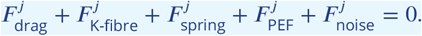

The drag force is taken as proportional to kinetochore velocity,

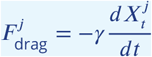

where *γ* is the effective drag coefficient. The K-fibre force, 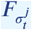, is dependent on the hidden microtubule polymerization state, and other forces are as described in the main text. Thus we find

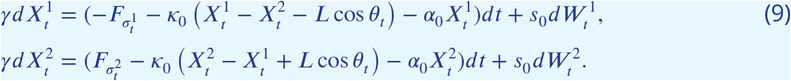

Dividing by the drag coefficient, *γ*, and redefining rescaled force parameters for the PEF, *α* = *α*_0_/*γ*, spring constant, *κ* = *κ*_0_/*γ*, noise magnitude, *s* = *s*_0_/*γ* and microtubule force parameters *ν*_+_ = *F*_+_/*γ, ν*_−_ = *F*_−_/*γ*, we obtain

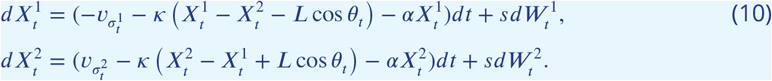

Due to this rescaling by the drag coefficient, which is hard to estimate, the K-fibre force parameters (*ν*_+_, *ν*_−_) that we infer in this work have units of speed [um/s], while the PEF parameter, *α*, and spring constant, *κ*, have units of [s^−1^].

## Footnotes

1 *q*_*AR*_ = 1 − *p*_*AR*_ is thus the probability of switching from the anaphase state, *A*, to the reversal state, *R*.

## References

Armond JW, Dale KL, Burroughs NJ, McAinsh AD, Vladimirou E. The dynamics of centromere motion through the metaphase-to-anaphase transition reveal a centromere separation order. BioRxiv. 2019; p. 582379.

Armond JW, Harry EF, McAinsh AD, Burroughs NJ. Inferring the forces controlling metaphase kinetochore oscillations by reverse engineering system dynamics. PLoS Computational Biology. 2015; 11(11):e1004607.

Armond JW, Vladimirou E, McAinsh AD, Burroughs NJ. KiT: a MATLAB package for kinetochore tracking. Bioinformatics. 2016; 32(12):1917–1919.

Auckland P, Clarke NI, Royle SJ, McAinsh AD. Congressing kinetochores progressively load Ska complexes to prevent force-dependent detachment. Journal of Cell Biology. 2017; 216(6):1623–1639.

Bakhoum SF, Genovese G, Compton DA. Deviant kinetochore microtubule dynamics underlie chromosomal instability. Current Biology. 2009; 19(22):1937–1942.

Blackwell R, Sweezy-Schindler O, Edelmaier C, Gergely ZR, Flynn PJ, Montes S, Crapo A, Doostan A, McIntosh JR, Glaser MA, Betterton MD. Contributions of microtubule dynamic instability and rotational diffusion to kinetochore capture. Biophysical Journal. 2017; 112(3):552–563.

Browning AP, Warne DJ, Burrage K, Baker RE, Simpson MJ. Identifiability analysis for stochastic differential equation models in systems biology. Journal of the Royal Society Interface. 2020; 17(173):20200652.

Burroughs NJ, Harry EF, McAinsh AD. Super-resolution kinetochore tracking reveals the mechanisms of human sister kinetochore directional switching. Elife. 2015; 4:e09500.

Carpenter B, Gelman A, Hoffman MD, Lee D, Goodrich B, Betancourt M, Brubaker MA, Guo J, Li P, Riddell A. Stan: a probabilistic programming language. Journal of Statistical Software. 2017; 76(1):1–32.

Chen BC, Legant WR, Wang K, Shao L, Milkie DE, Davidson MW, Janetopoulos C, Wu XS, Hammer JA, Liu Z, et al. Lattice light-sheet microscopy: imaging molecules to embryos at high spatiotemporal resolution. Science. 2014; 346(6208).

Civelekoglu-Scholey G, Cimini D. Modelling chromosome dynamics in mitosis: A historical perspective on models of metaphase and anaphase in eukaryotic cells. Interface Focus. 2014; 4(3).

Civelekoglu-Scholey G, He B, Shen M, Wan X, Roscioli E, Bowden B, Cimini D. Dynamic bonds and polar ejection force distribution explain kinetochore oscillations in PtK1 cells. Journal of Cell Biology. 2013; 201(4):577–593.

Cuylen S, Blaukopf C, Politi AZ, Müller-Reichert T, Neumann B, Poser I, Ellenberg J, Hyman AA, Gerlich DW. Ki-67 acts as a biological surfactant to disperse mitotic chromosomes. Nature. 2016; 535(7611):308–312.

Drpic D, Almeida AC, Aguiar P, Renda F, Damas J, Lewin HA, Larkin DM, Khodjakov A, Maiato H. Chromosome segregation is biased by kinetochore size. Current Biology. 2018; 28(9):1344–1356.

Dumont M, Gamba R, Gestraud P, Klaasen S, Worrall JT, De Vries SG, Boudreau V, Salinas-Luypaert C, Maddox PS, Lens SM, et al. Human chromosome-specific aneuploidy is influenced by DNA-dependent centromeric features. The EMBO journal. 2020; 39(2):e102924.

Elting MW, Prakash M, Udy DB, Dumont S. Mapping load-bearing in the mammalian spindle reveals local kinetochore fiber anchorage that provides mechanical isolation and redundancy. Current Biology. 2017; 27(14):2112–2122.

Elting MW, Suresh P, Dumont S. The spindle: Integrating architecture and mechanics across scales. Trends in Cell Biology. 2018; 28(11):896–910.

Ferrandiz N, Downie L, Starling GP, Royle SJ. Endomembranes promote chromosome missegregation by ensheathing misaligned chromosomes. bioRxiv. 2021;.

Gelman A, Carlin JB, Stern HS, Dunson DB, Vehtari A, Rubin DB. Bayesian Data Analysis, vol. 2. CRC press Boca Raton, FL; 2014.

Gelman A, Rubin DB. Inference from iterative simulation using multiple sequences. Statistical Science. 1992; 7(4):457–472.

Gregan J, Polakova S, Zhang L, Tolić-Nørrelykke IM, Cimini D. Merotelic kinetochore attachment: causes and effects. Trends in Cell Biology. 2011; 21(6):374–381.

Harasymiw LA, Tank D, McClellan M, Panigrahy N, Gardner MK. Centromere mechanical maturation during mammalian cell mitosis. Nature communications. 2019; 10(1):1–21.

Hauf S, Waizenegger IC, Peters JM. Cohesin cleavage by separase required for anaphase and cytokinesis in human cells. Science. 2001; 293(5533):1320–1323.

Hawkes AG. Spectra of some self-exciting and mutually exciting point processes. Biometrika. 1971; 58(1):83–90.

Hill TL. Theoretical problems related to the attachment of microtubules to kinetochores. Cell Biology. 1985; 82:4404–4408.

Hines KE, Middendorf TR, Aldrich RW. Determination of parameter identifiability in nonlinear biophysical models: A Bayesian approach. The Journal of General Physiology. 2014; 143(3):401–416.

Hoffman MD, Gelman A. The No-U-Turn sampler: adaptively setting path lengths in Hamiltonian Monte Carlo. Journal of Machine Learning Research. 2014; 15(1):1593–1623.

Holt LJ, Krutchinsky AN, Morgan DO. Positive feedback sharpens the anaphase switch. Nature. 2008; 454(7202):353–357.

Iemura K, Natsume T, Maehara K, Kanemaki MT, Tanaka K. Chromosome oscillation promotes Aurora A– dependent Hec1 phosphorylation and mitotic fidelity. Journal of Cell Biology. 2021; 220(7):e202006116.

Jaqaman K, King EM, Amaro AC, Winter JR, Dorn JF, Elliott HL, Mchedlishvili N, McClelland SE, Porter IM, Posch M, et al. Kinetochore alignment within the metaphase plate is regulated by centromere stiffness and microtubule depolymerases. Journal of Cell Biology. 2010; 188(5):665–679.

Joglekar AP, Hunt AJ. A simple, mechanistic model for directional instability during mitotic chromosome movements. Biophysical Journal. 2002; 83(1):42–58.

Kajtez J, Solomatina A, Novak M, Polak B, Vukušić K, Rüdiger J, Cojoc G, Milas A, Šestak IŠ, Risteski P, et al. Overlap microtubules link sister k-fibres and balance the forces on bi-oriented kinetochores. Nature Communications. 2016; 7(1):1–11.

Ke K, Cheng J, Hunt AJ. The distribution of polar ejection forces determines the amplitude of chromosome directional instability. Current Biology. 2009; 19(10):807–815.

Loeffer C, Flaxman S. Is gun violence contagious? A spatiotemporal test. Journal of Quantitative Criminology. 2018; 34(4):999–1017.

McIntosh JR, Landis SC. The distribution of spindle microtubules during mitosis in cultured human cells. The Journal of Cell Biology. 1971; 49(2):468–497.

Miles CE, Zhu J, Mogilner A. Mechanical torque promotes bipolarity of the mitotic spindle through multicentrosomal clustering. bioRxiv. 2021; doi: 10.1101/2021.11.17.469054.

Mogilner A, Wollman R, Civelekoglu-Scholey G, Scholey J. Modeling mitosis. Trends in Cell Biology. 2006; 16(2):88–96.

Musacchio A. Spindle assembly checkpoint: The third decade. Philosophical Transactions of the Royal Society B: Biological Sciences. 2011; 366(1584):3595–3604.

Neal RM. MCMC using Hamiltonian dynamics. Handbook of Markov Chain Monte Carlo. 2011; 2(11):2.

Novak M, Polak B, Simunić J, Boban Z, Kuzmić B, Thomae AW, Tolić IM, Pavin N. The mitotic spindle is chiral due to torques within microtubule bundles. Nature Communications. 2018; 9(1):1–10.

Pargett M, Umulis DM. Quantitative model analysis with diverse biological data : Applications in developmental pattern formation Labeled structures. Methods. 2013; 62(1):56–67.

Paul R, Wollman R, Silkworth WT, Nardi IK, Cimini D, Mogilner A. Computer simulations predict that chromosome movements and rotations accelerate mitotic spindle assembly without compromising accuracy. . 2009; 106(37):15708–15713.

Pavin N, Tolić IM. Mechanobiology of the mitotic spindle. Developmental Cell. 2020;.

Polak B, Risteski P, Lesjak S, Tolic IM. PRC 1-labeled microtubule bundles and kinetochore pairs show one-toone association in metaphase. EMBO reports. 2017; 18(2):217–230.

Rabiner LR. A tutorial on hidden Markov models and selected applications in speech recognition. Proceedings of the IEEE. 1989; 77(2):257–286.

Rago F, Cheeseman IM. The functions and consequences of force at kinetochores. Journal of Cell Biology. 2013; 200(5):557–565.

Reinhart A. A review of self-exciting spatio-temporal point processes and their applications. Statistical Science. 2018; 33(3):299–318.

Roscioli E, Germanova TE, Smith CA, Embacher PA, Erent M, Thompson AI, Burroughs NJ, McAinsh AD. Ensemble-level organization of human kinetochores and evidence for distinct tension and attachment sensors. Cell Reports. 2020; 31(4):107535.

Sen O, Harrison JU, Burroughs NJ, McAinsh AD. Kinetochore life histories reveal an Aurora-B-dependent error correction mechanism in anaphase. Developmental Cell. 2021;.

Shimamoto Y, Maeda YT, Ishiwata S, Libchaber AJ, Kapoor TM. Insights into the micromechanical properties of the metaphase spindle. Cell. 2011; 145(7):1062–1074.

Simunić J, Tolić IM. Mitotic spindle assembly: building the bridge between sister K-fibers. Trends in Biochemical Sciences. 2016; 41(10):824–833.

Skibbens RV, Skeen VP, Salmon ED. Directional Instability of Kinetochore Motility during Chromosome Congression and Segregation in Mitotic Newt Lung Cells : A Push-Pull Mechanism. Journal of Cell Biology. 1993; 122(4).

Smith CA, McAinsh AD, Burroughs NJ. Human kinetochores are swivel joints that mediate microtubule attachments. Elife. 2016; 5:e16159.

Stephens AD, Haggerty RA, Vasquez PA, Vicci L, Snider CE, Shi F, Quammen C, Mullins C, Haase J, Taylor RM, et al. Pericentric chromatin loops function as a nonlinear spring in mitotic force balance. Journal of Cell Biology. 2013; 200(6):757–772.

Su KC, Barry Z, Schweizer N, Maiato H, Bathe M, Cheeseman IM. A regulatory switch alters chromosome motions at the metaphase-to-anaphase transition. Cell Reports. 2016; 17(7):1728–1738.

Tolić IM. Mitotic spindle: kinetochore fibers hold on tight to interpolar bundles. European Biophysics Journal. 2018; 47(3):191–203.

Vázquez-Novelle MD, Sansregret L, Dick AE, Smith CA, McAinsh AD, Gerlich DW, Petronczki M. Cdk1 inactivation terminates mitotic checkpoint surveillance and stabilizes kinetochore attachments in anaphase. Current Biology. 2014; 24(6):638–645.

Vehtari A, Gelman A, Simpson D, Carpenter B, Bürkner PC. Rank-normalization, folding, and localization: An improved ?^ for assessing convergence of MCMC. Bayesian Analysis. 2021; 1(1):1–28.

Vladimirou E, Mchedlishvili N, Gasic I, Armond JW, Samora CP, Meraldi P, McAinsh AD. Nonautonomous movement of chromosomes in mitosis. Developmental Cell. 2013; 27(1):60–71.

Wan X, Cimini D, Cameron LA, Salmon E. The coupling between sister kinetochore directional instability and oscillations in centromere stretch in metaphase PtK1 cells. Molecular Biology of the Cell. 2012; 23(6):1035–1046.

Willms AR, Kitanov PM, Langford WF. Huygens’ clocks revisited. Royal Society open science. 2017; 4(9):170777.

Worrall JT, Tamura N, Mazzagatti A, Shaikh N, van Lingen T, Bakker B, Spierings DCJ, Vladimirou E, Foijer F, McClelland SE. Non-random mis-segregation of human chromosomes. Cell Reports. 2018; 23(11):3366–3380.

Zaytsev AV, Grishchuk EL. Basic mechanism for biorientation of mitotic chromosomes is provided by the kinetochore geometry and indiscriminate turnover of kinetochore microtubules. Molecular biology of the cell. 2015; 26(22):3985–3998.

